# How to build a fast and highly sensitive sound detector that remains robust to temperature shifts

**DOI:** 10.1101/673186

**Authors:** Minghui Chen, Henrique von Gersdorff

## Abstract

Frogs must have sharp hearing abilities during the warm summer months to successfully find mating partners. This study aims to understand how frog hair cell ribbon-type synapses preserve both sensitivity and temporal precision during temperature changes. We performed *in vitro* patch-clamp recordings of hair cells and their afferent fibers in bullfrog amphibian papillae under room (23-25°C) and high (30-33°C) temperature. Afferent fibers exhibited a wide heterogeneity in membrane input resistance (R_in_) from 100 MΩ to 1000 MΩ, which may contribute to variations in spike threshold and firing frequency. At higher temperatures, most fibers increased their frequency of action potential firing due to an increase in spontaneous EPSC frequencies. Hair cell resting membrane potential (V_rest_) remained surprisingly stable during temperature increases, although both inward Ca^2+^ current and outward K^+^ current increased in amplitude. This increase in Ca^2+^ current may explain the higher spontaneous EPSC frequencies. The larger “leak currents” at V_rest_ lowered R_in_ and produced higher electrical resonant frequencies. However, lower R_in_ should decrease sensitivity to sound detection via smaller receptor potentials. Using membrane capacitance measurements, we suggest that hair cells can partially compensate for this reduced sensitivity by increasing exocytosis efficiency and the size of the readily releasable pool of synaptic vesicles. Furthermore, paired recordings of hair cells and their afferent fibers showed that synaptic delays become shorter and multivesicular release becomes more synchronous at higher temperatures, which should improve temporal precision. Altogether, our results explain many previous *in vivo* observations on the temperature dependence of spikes in auditory nerves.

**Significance Statement:** The vertebrate inner ear detects and transmits auditory information over a broad dynamic range of sound frequency and intensity. It achieves remarkable sensitivity to soft sounds and precise frequency selectivity. How does the ear of cold-blooded vertebrates maintain its performance level as temperature changes? More specifically, how does the hair cell to afferent fiber synapse in bullfrog amphibian papilla adjust to a wide range of physiological temperatures without losing its sensitivity and temporal fidelity to sound signals? This study uses *in vitro* experiments to reveal the biophysical mechanisms that explain many observations made from *in vivo* auditory nerve fiber recordings. We find that higher temperature facilitates vesicle exocytosis and electrical tuning to higher sound frequencies, which benefits sensitivity and selectivity.

## Introduction

Bullfrogs that jump into a cold pond must adapt quickly to a sudden temperature shift. Indeed, frogs adapt to a wide range of physiological body temperatures (Lillywhite, 1970). In summer, male frogs gather together and chorus to attract female frogs (Capranica, 1965). To successfully find mating partners their hearing abilities must be sharp during the warm summer months. Bullfrog vocalization is composed of low- to mid-frequency sounds (200-2000 Hz; Capranica, 1965; Smotherman and Narins, 2000; Simmons, 2004). In addition, hearing is also critical for hunting insects and for territorial behaviors (Emlen, 1968; Wiewandt, 1969). Insects flap their wings at frequencies ranging from 330 Hz (house flies) to 600 Hz (mosquitos), and can even reach 1040 Hz (small biting midges). All these sound frequencies are detected and transduced by one frog hearing organ, the amphibian papilla (van Dijk et al., 2011). Accordingly, recordings of spikes in bullfrog amphibian papilla auditory nerve fibers display tuning curves with greatest sensitivity and best frequencies (or lowest sound level thresholds) around 500-650 Hz (Feng et al., 1975; Heffner and Heffner, 2007).

Hair cells transduce sound vibrations into graded electrical signals, which are then sent to the brain via all-or-none action potential spikes in the afferent fibers. At higher temperatures, *in vivo* single afferent fiber recordings have revealed an increase in spontaneous spike rates, a decrease in sound intensity threshold, a reduced latency of response to sound and higher vector strength (or better phase-locking precision; Stiebler and Narins, 1990; van Dijk et al., 1990). This indicates that the hearing organ of frogs transmit more sound information with higher sensitivity, shorter reaction times and greater temporal precision at higher temperatures. What are the cellular and synaptic mechanisms that explain these *in vivo* observations?

Hair cells detect and transduce three aspects of sound: intensity, phase and frequency. Information on the rapid onset and offset of sound transients must also be faithfully transmitted to the auditory nerves. Indeed, hair cells express ion channels with some of the fastest activation and deactivation kinetics (Engel, 2008; Heil and Peterson, 2017; Pangrsic et al., 2018). Sound signals are conveyed via transduction currents (I) mediated by K^+^ influx at the stereocilia bundles, resulting in graded receptor membrane potential (V_m_) changes. The detection of low-level sounds is facilitated if hair cells have a large input resistance (R_in_), given that V_m_ = R_in_ × I. However, phase-locking to higher frequency sounds with fine temporal precision requires shorter membrane time constants (τ_m_ = R_in_ × C_m_, where C_m_ is the hair cell membrane capacitance), which requires a small R_in_. How does the hair cell cope with these conflicting demands on its biophysical properties? Does hair cell R_in_ decrease when temperature increases, as observed in other bullfrog neurons (Santin et al., 2013)? If so, how do auditory hair cells and their synapses compensate for temperature-dependent changes in R_in_ to maintain both sound sensitivity and temporal fidelity?

To answer these questions, we performed *in vitro* voltage-clamp and current-clamp recordings from single hair cells and their afferent fibers in bullfrog amphibian papillae under both room (23-25°C) and high (30-33°C) temperature. Our results suggest that larger amplitudes and faster Ca^2+^ and K^+^ current kinetics lead to higher hair cell intrinsic electrical resonance frequencies, whereas shorter synaptic delays, more synchronous multivesicular release, and decreased R_in_ at high temperature contributes to more precise phase locking to sound signals. Moreover, we propose that hair cells compensate for lower R_in_ at high temperature by increasing the size of the readily releasable pool of vesicles and the efficiency of exocytosis, resulting in an enhancement of sound sensitivity.

## Materials and Methods

Adult bullfrogs (*Rana catesbeiana*; Rana Ranch, Twin Falls, ID) of both sexes were used for experiments. Bullfrogs were sedated in 7-10°C water bath for about 5-10 min, anesthetized by isoflurane (0.025 ml/g body weight) absorbed through the skin, and then double-pitched and decapitated. Protocols were approved by the Institutional Animal Care and Use Committee of Oregon Health and Science University.

Amphibian papillae were carefully dissected and the connection of hair cells and afferent fibers was exposed following the protocols described previously (Keen and Hudspeth, 2006; Li et al., 2009). The acutely split-open tissue preparation was placed in a recording chamber perfused with artificial perilymph containing (in mM): 95 NaCl, 2 KCl, 2 CaCl_2_, 1 MgCl_2_, 25 NaHCO_3_, 3 glucose, 1 creatine and 1 sodium pyruvate (pH 7.30, osmolarity 235 mOsm) at 1-2 ml/min, bubbling with 95% O_2_ and 5% CO_2_. Temperature was adjusted by heating the bath perfusion with a temperature controller (Warner Instruments), which changed bath temperature between room (23-25°C) and high (30-33°C) temperatures. Temperature was measured continuously by a miniature thermistor placed close to the perfused amphibian papillae preparation. Reagents were obtained from Millipore-Sigma (St. Louis, MO).

### Electrophysiology

Semi-intact preparations of hair cells and their connecting afferent fibers were placed on an upright microscope with a 60× water-immersion objective (Olympus BX51WI) and digital CCD camera (QImaging). Recording pipettes were pulled on a PP-830 vertical puller (Narishige) from borosilicate glass pipettes (1B150F-4, World Precision Instruments). Patch pipettes were pulled to resistances of 5-7 MΩ for hair cells. Pipettes were filled with the intracellular solution containing the following (in mM): 77 Cs-gluconate, 20 CsCl, 1 MgCl_2_, 10 TEA-Cl, 10 HEPES, 2 EGTA, 3 Mg-ATP, 1 Na-GTP, and 5 Na_2_-phosphocreatine, adjusted to pH 7.3 with CsOH. We used 2 mM EGTA as the mobile internal Ca^2+^ buffer of hair cells (Frank et al., 2009; Johnson et al., 2017). The pH-independent Ca^2+^ buffer BAPTA (2 mM) and another pH buffer MOPS (10 mM) were also used in some experiments (Fig. 9). To measure resting membrane potential (Fig. 5 A), outward potassium current (Fig. 5 B and D), membrane input resistance (Fig. 6) and resonant frequency (Fig. 7), we used a more physiological K^+^-based internal solution containing the following (in mM): 77 K-gluconate, 30 KCl, 1 MgCl_2_, 10 HEPES, 2 EGTA, 3 Mg-ATP, 1 Na-GTP, and 5 Na_2_-phosphocreatine, adjusted to pH 7.3 with KOH. Whole-cell patch recordings were performed with an EPC-10/2 (HEKA) patch-clamp amplifier controlled by PatchMaster software (HEKA). In voltage clamp, hair cells were held at a membrane potential of −90 mV. Membrane potentials were corrected for a liquid junction potential of 10 mV. In current clamp, hair cells were held at zero current. Whole-cell calcium currents (I_Ca_) were leak subtracted using a P/4 protocol. The uncompensated series resistances (R_s_) in whole-cell recordings were 10.5 ± 0.4 MΩ for hair cells (n = 36). To preserve a more intact hair cell intracellular milieu we also performed perforated-patch recordings using an internal solution with gramicidin (40-50 μg/ml; Fig. 5A and 6) plus a fluorescent dye (Alexa Fluor 488 Hydrazide; ThermoFisher Scientific). Hair cells containing dye in their cytoplasm were excluded from analysis. The uncompensated average R_s_ in perforated-patch recordings was 40.1 ± 1.3 MΩ (n = 9).

We used whole-cell membrane capacitance measurement techniques to measure the increase in membrane capacitance (C_m_) accompanying vesicle fusion. Patch pipettes were coated with dental wax to minimize their stray capacitance. The C_m_ from hair cells were measured under whole-cell voltage-clamp conditions using the “Sine + DC” method (Lindau and Neher, 1988; Moser and Beutner, 2000). A 2 kHz sinusoidal command voltage of 50 mV peak-to-peak magnitude was superposed on the hair cell holding potential of −90 mV. The resulting current response was used to calculate C_m_ via a software emulator of a lock-in amplifier (Heka EPC-10; Gilles, 2000). The increase of C_m_(ΔC_m_) evoked by membrane depolarization was measured as ΔC_m_ = C_m_(response) − C_m_(resting). Here C_m_(resting) and C_m_(response) were obtained by averaging capacitance data points before and after the depolarizing steps using Igor Pro 6.0 (WaveMetrics, OR) software. We excluded recordings in which the uncompensated series resistance was above 15 MΩ or the holding current was larger than 100 pA.

Afferent fibers were patched with 8-10 MΩ patch pipette filled with the K^+^-based internal solution, as described above. Membrane potentials were corrected for a liquid junction potential of 10 mV. In voltage clamp, afferent fibers were held at a membrane potential of −90 mV. Spontaneous excitatory post-synaptic current (EPSCs) were recorded from the voltage-clamped afferent fibers. Spontaneous excitatory post-synaptic potentials (EPSPs) were recorded in current-clamp mode from afferent fibers with zero current injection. The uncompensated R_s_ in whole-cell recordings of afferent fibers were 27.9 ± 3.2 MΩ (n = 8). An excessively high series resistance can reduce and filter large EPSC events because of significant voltage-clamp errors (Li et al., 2009). Therefore, we excluded afferent fibers with uncompensated series resistance above 50 MΩ from analysis and we electronically compensated afferent fiber whole-cell recordings up to 35% depending on the uncompensated series resistance to maintain a constant series resistance throughout the recordings.

### Data analysis

Statistical analysis was performed using Prism 6 (GraphPad Software) and Excel (Microsoft). Results are presented as mean ± SEM (n = number of cells). If not specified, statistical significance was determined using paired Student’s *t* test. We chose *p* < 0.05 to be the criterion for statistical significance. Data analyses and curve fitting were performed using Igor Pro 6.0. The threshold of an action potential was defined as membrane potential (V_m_) when the slope of V_m_ versus time started to increase compared to that of preceding EPSP. Our estimation was consistent with the threshold estimated by phase plots, which plot membrane potential slope (dV_m_/dt) versus V_m_ (Yang et al., 2016). Biophysical properties of spontaneous EPSCs and EPSPs (i.e., amplitude, frequency, rise time and τ_decay_) were analyzed using with a customized protocol written by Dr. Owen Gross. The rise time was calculated as the interval between 10% and 90% of the peak amplitude relative to baseline. The τ_decay_ of averaged EPSCs or EPSPs was estimated by a single exponential fit using the equation:

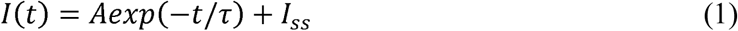

where I(t) is the current as a function of time, A is the amplitude at time 0, τ is the time constant, and I_ss_ is the steady-state current amplitude. The temperature coefficient (Q_10_) is the coefficient by which a quantity increases after a change of 10°C. Q_10_ values for amplitude, frequency, rise time and time constant of decay as well as ΔC_m_ were calculated using the equation:

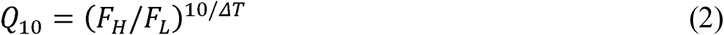

where F_L_ and F_H_ are factors of interest measured at room (23-25°C) and high (30-33°C) temperature, respectively, and ΔT is the absolute value of the temperature difference.

## Results

### Most afferent fibers fire more spikes at higher temperature

To determine the temperature dependence of spontaneous spikes at hair cell afferent fibers, we recorded spontaneous potential changes in afferent fibers using whole-cell current clamp with zero current injection. At 23.7 ± 0.2°C, the frequency of spikes was 2.0 ± 0.64 Hz, amplitude was 51.9 ± 6.8 mV, threshold was −51.0 ± 2.0 mV and resting membrane potential (V_rest_) of afferent fibers was −66.9 ± 2.1 mV (n = 18). Among these fibers, seven of them displayed big spikes that overshot 0 mV with higher threshold (Fig. 1A1). The amplitude of these big spikes was 85.7 ± 5.7 mV (n = 6), whereas the amplitude of spikes that did not overshoot was 35.0 ± 4.8 mV (n = 12, *p* < 0.0001). The threshold of the big spikes was −44.1 ± 2.4 mV (n = 6), whereas the threshold of small spikes was −54.8 ± 2.1 mV (n = 12, *p* = 0.0063, unpaired *t* test). There was no statistical difference between these two groups in frequency (overshooting: 2.02 ± 0.56 Hz versus non-overshooting: 1.99 ± 0.94 Hz, *p* = 0.98, unpaired *t* test) and V_rest_ (overshooting: −64.1 ± 4.6 mV versus non-overshooting: −68.3 ± 2.3 mV, *p* = 0.37).

**Fig. 1.**
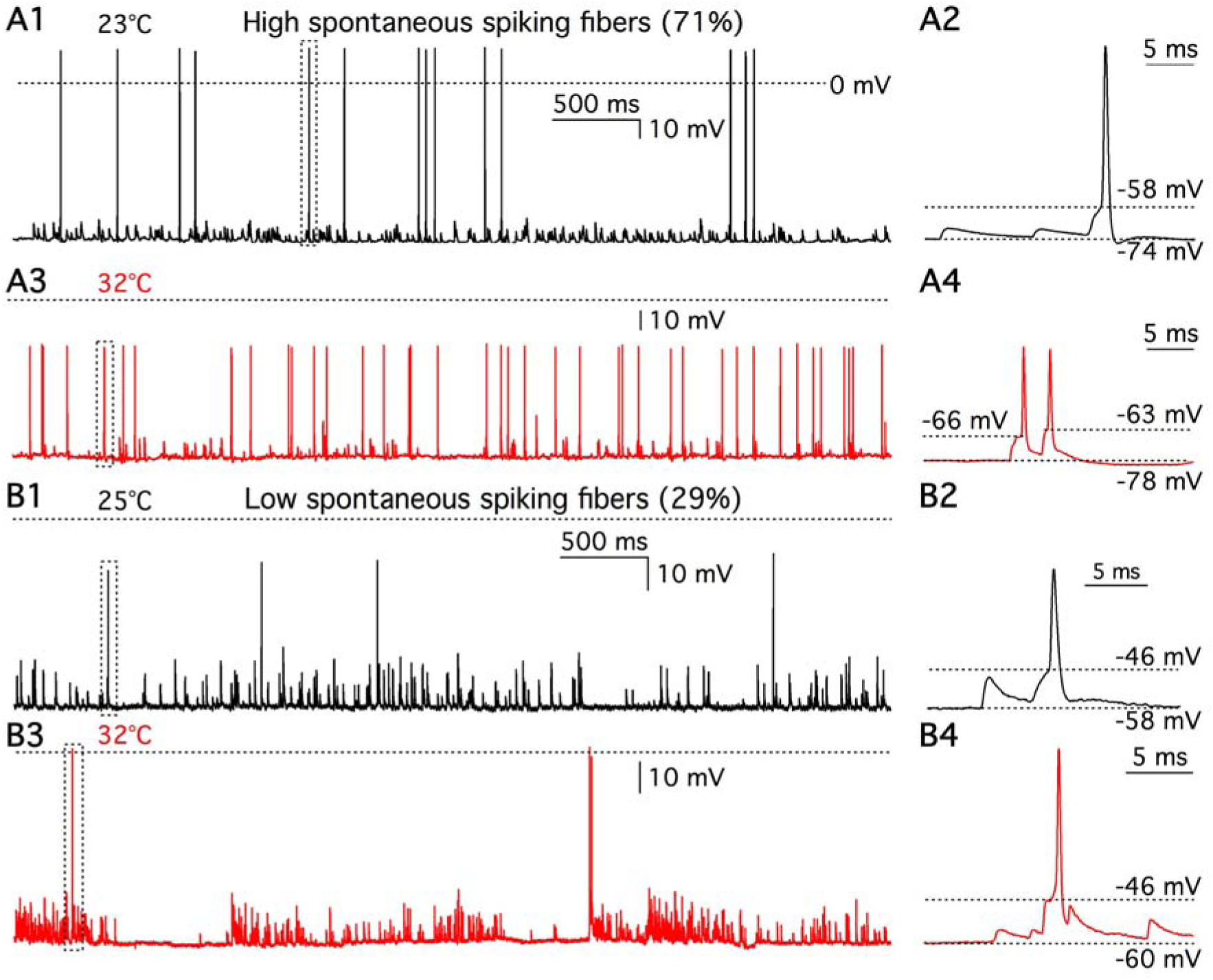
Heterogeneity in the temperature dependence of afferent fiber spikes. (A1) Whole cell current clamp recordings of afferent fiber with zero current injection at 23°C. Action potential (AP) spikes and EPSPs can be clearly distinguished. The dash line indicates 0 mV. (A2) The spike shown in a dash line box in (A1) was expanded in time scale. Resting membrane potential (V_rest_) was −74 mV and AP threshold was −58 mV. (A3) Spontaneous spikes from the same afferent fiber shown in (A1) were recorded at 32°C. (A4) Spikes shown in a dash line box in (A3) are expanded in time scale. V_rest_ was −78 mV. Thresholds of the 1^st^ and the 2^nd^ AP were −66 mV and −63 mV, respectively. Four out of six afferent fibers (68%) fired more spikes at high temperature. (B1) Spontaneous spikes were recorded from another afferent fiber at 25°C. The dash line indicates 0 mV. (B2) The spike shown in a dash line box in (B1) was expanded in time scale. V_rest_ was −59 mV and AP threshold was −47 mV. (B3) Spontaneous spikes from the same fiber shown in (B1) were recorded at 32°C. (B4) The spike shown in a dash line box in (B3) was expanded in time scale. V_rest_ was −60 mV and AP threshold was −46 mV. Two out of six afferent fibers (32%) fired less spikes at high temperature.

For seven out of these 18 fibers, we successfully recorded spikes at both room and high temperature. Five of them fired more spikes at high temperature (Table 1, Fig. 1A1-A4). We called these fibers “thermal positive fibers”. The temperature coefficient (Q_10_) of spontaneous spiking rate was 4.0 ± 1.2 (n = 5), which was similar to that found in rat fibers (median Q_10_ = 6.6; Wu et al, 2016). Although threshold for firing an action potential at high temperature (−58.6 mV) was lower than that at room temperature (−53.9 mV, n = 5, *p* = 0.0158), spike probability rates remain the same throughout temperature change: at room temperature 25.8% of EPSPs evoked a spike and at high temperature 25.4% of EPSPs evoked a spike (n = 5, *p* = 0.97). A decrease in afferent fiber membrane input resistance at high temperature results in faster EPSP decay (see Fig. 4) making it more difficult for them to summate, which counteracts the potential increase in spike probability caused by lower thresholds. By contrast, in mammalian cochlear spiral ganglion neurons (SGN), 80-97% of spontaneous EPSPs evoke a spike under room temperature (Rutherford et al, 2012). The low spike failure rate of EPSPs in mammalian fibers may result from the smaller surface area and high input resistance of the mammalian postsynaptic bouton (Rutherford et al, 2012). In bullfrog amphibian papilla, each hair cell connects with 3 to 6 large claw-like afferent fiber endings that receive multiple ribbon synapses (Graydon et al., 2014), while each mammalian afferent fiber only connects with just one single synaptic ribbon (Liberman, 1980; Rutherford, 2015).

**Table 1.**
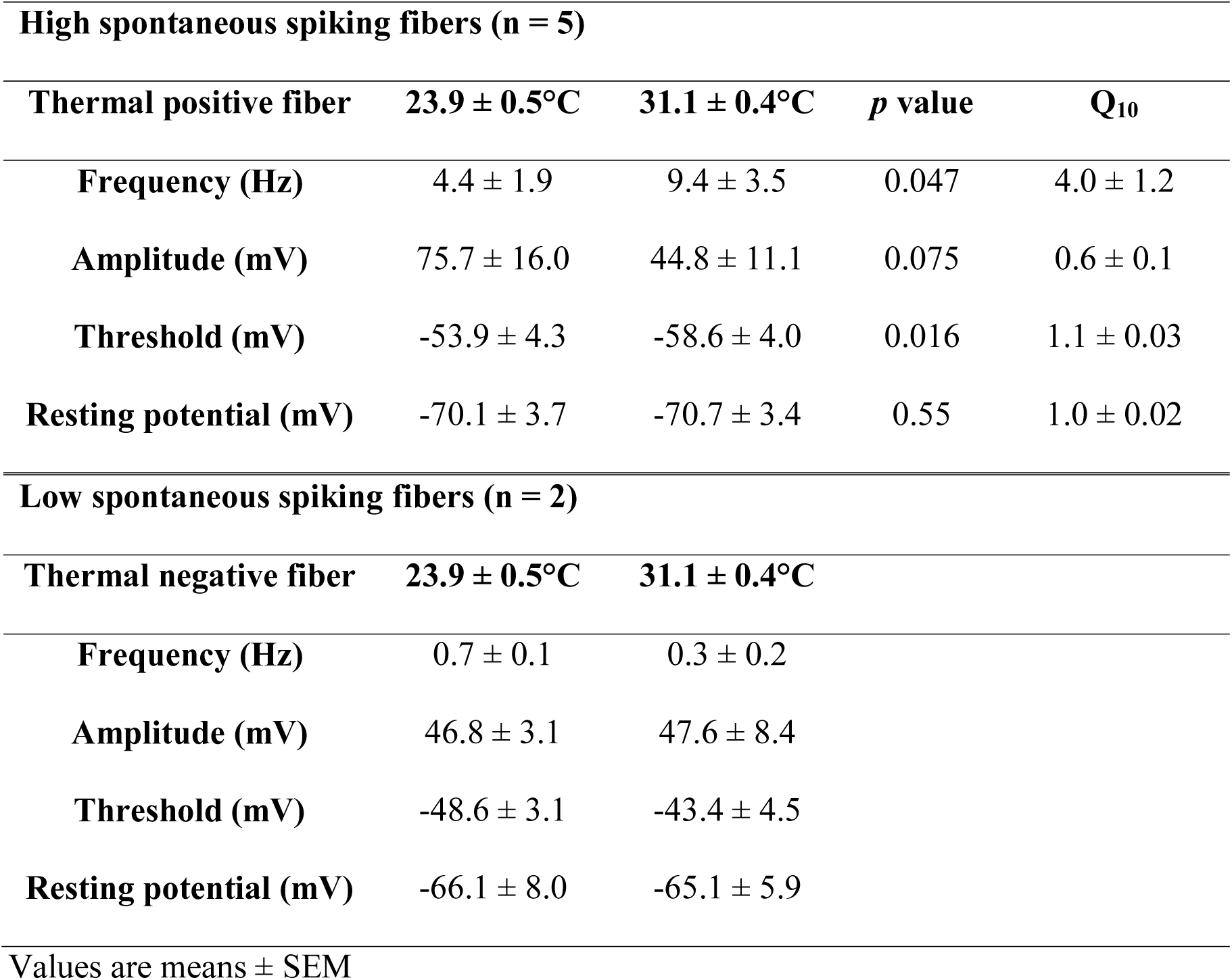
Temperature dependence of afferent fiber spikes

| **High spontaneous spiking fibers (n = 5)** |  |  |  |  |
| --- | --- | --- | --- | --- |
| **Thermal positive fiber** | **23.9 ± 0.5°C** | **31.1 ± 0.4°C** | ***p* value** | **Q<sub>10</sub>** |
| **Frequency (Hz)** | 4.4 ± 1.9 | 9.4 ± 3.5 | 0.047 | 4.0 ± 1.2 |
| **Amplitude (mV)** | 75.7 ± 16.0 | 44.8 ± 11.1 | 0.075 | 0.6 ± 0.1 |
| **Threshold (mV)** | -53.9 ± 4.3 | -58.6 ± 4.0 | 0.016 | 1.1 ± 0.03 |
| **Resting potential (mV)** | -70.1 ± 3.7 | -70.7 ± 3.4 | 0.55 | 1.0 ± 0.02 |
| **Low spontaneous spiking fibers (n = 2)** |  |  |  |  |
| **Thermal negative fiber** | **23.9 ± 0.5°C** | **31.1 ± 0.4°C** |  |  |
| **Frequency (Hz)** | 0.7 ± 0.1 | 0.3 ± 0.2 |  |  |
| **Amplitude (mV)** | 46.8 ± 3.1 | 47.6 ± 8.4 |  |  |
| **Threshold (mV)** | -48.6 ± 3.1 | -43.4 ± 4.5 |  |  |
| **Resting potential (mV)** | -66.1 ± 8.0 | -65.1 ± 5.9 |  |  |

Values are means ± SEM

The other two fibers fired less spikes at high temperature (Table 1, Fig. 1B1-B4) and we called these “thermal negative fibers”. The thermal negative fibers had low spontaneous spiking rate (< 1Hz, Table 1). In rat auditory nerve, spontaneous spiking rate increases during maturation of the cochlear from 3.87 Hz at P15-P17 to 12.85 Hz at P29-P32 (Wu et al, 2016). It is possible that some low frequency spiking fibers in the frog may receive synaptic release from some immature hair cells at the edge of the amphibian papilla (Lewis and Li, 1975).

### Frequencies of spontaneous EPSCs and EPSPs are enhanced at high temperature

To examination how temperature affects hair cell spontaneous release, we recorded spontaneous EPSCs from afferent fibers without voltage clamping their presynaptic hair cells under room (black traces) and high (red traces) temperature (Fig. 2A1 and B1). The frequency of the EPSCs increased from 78.2 ± 9.5 Hz at room temperature to 115.2 ± 9.7 Hz at high temperature (n = 18, *p* = 0.0097, Table 2). The EPSC amplitude distribution of an afferent fiber in Fig. 2A2 also shows that more spontaneous EPSCs occurred under high temperature. The EPSC amplitude distribution was fit well with a Gaussian function (Fig. 2A2: R^2^ = 0.96 at 22°C; R^2^ = 0.93 at 32°C). The peaks of Gaussian fits fell at 93.5 pA under 22°C and at 89.7 pA under 32°C (Fig. 2A2), suggesting the average EPSC amplitude did not increase at high temperature. In Fig. 2A3, average EPSCs were normalized to their peak values and a single exponential function was used to fit the decay phase of EPSCs and determine the time constant of decay (τ_decay_). Surprisingly, the 67% of afferent fibers (12 out of 18) did not show an increase in EPSC amplitude at high temperature and the average of EPSC amplitude remained the same as temperature changed (*p* = 0.61, Table 2). This contrasts with results from the calyx of Held synapse (Kushmerick et al., 2006; Postlethwaite et al, 2007). The 10-90% rise time and τ_decay_ of spontaneous EPSCs were 223 ± 13 μs and 617 ± 36 μs (n =18, Table 2) at room temperature, which are faster than that measured in immature rat afferent fibers (Yi et al, 2010) but similar to that measured in the afferent fibers of adult turtle (Schnee et al, 2013) and hearing rats (Grant et al., 2010). Elevating temperature decreased both 10-90% rise time and τ_decay_ of spontaneous EPSCs with Q_10_ values of 1.7 and 1.8, respectively (Table 2). This is consistent with our previous findings showing that decreasing temperature from 25°C to 15°C slows down activation and decay of spontaneous EPSCs in bullfrog afferent fibers (Li et al., 2009). Similarly, in rat AII amacrine cells, increasing temperature from 26°C to 34°C decreases both rise time and τ_decay_ of mEPSCs with Q_10_ values of 1.3 and 1.5, respectively (Veruki et al., 2003). Calculating the integrals of spontaneous EPSCs, we found that the postsynaptic charge transfer was decreased at high temperature (Table 2), which may be due to faster activation and deactivation (Fig. 2A3 and B3) of postsynaptic AMPA receptors and faster glutamate transporter activity at high temperature (Auger and Attwell, 2000).

**Fig. 2.**
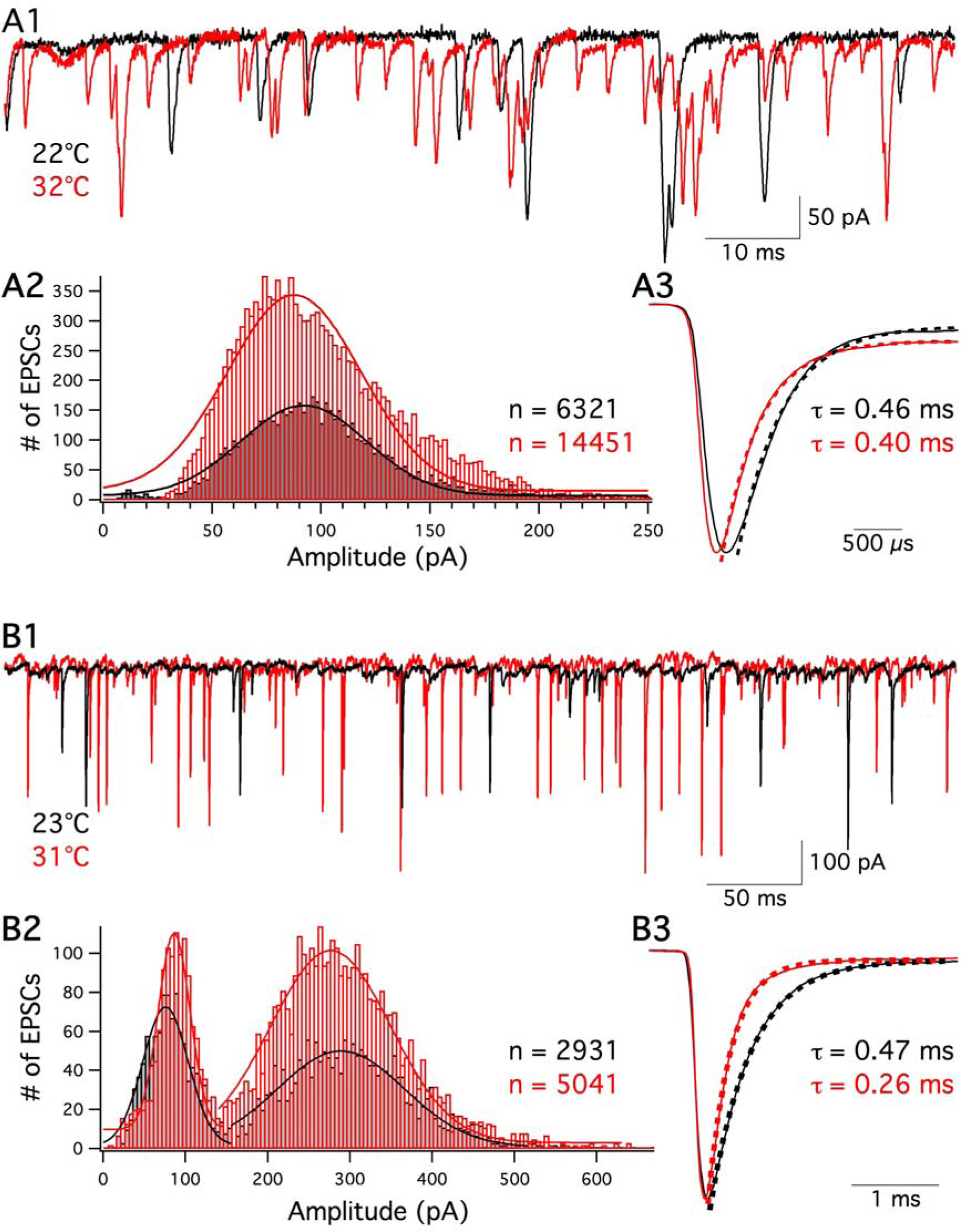
Spontaneous EPSC frequency increases at high temperature. (A1) Spontaneous EPSCs were recorded from an afferent fiber, which was voltage clamped at −90 mV at 22°C (black) and 32°C (red). (A2) A total of 6321 and 14451 EPSCs were obtained at 22°C (black) and 32°C (red) respectively. EPSC amplitude distributions were fit with a Gaussian function. (A3) Average of EPSCs recorded at 22°C (black) and 32°C (red) were normalized to their peak value. Single-exponential fits to the decay phase of averaged EPSCs showed that τ_decay_ were 0.46 ms at 22°C (black dash line) and 0.40 ms at 32°C (red dash line), respectively. (B1) Spontaneous EPSCs were recorded from another afferent fiber at 23°C (black) and 31°C (red). (B2) A total of 2931 and 5041 EPSCs were obtained at 23°C (black) and 31°C (red), respectively. Two peaks are present on the amplitude distribution. EPSCs were divided into two groups according to their size: small EPSCs with amplitude smaller than 150 pA and large EPSCs with amplitude larger than 150 pA. The EPSC amplitude distributions in each group were fit with a Gaussian function. (B3) Average of EPSCs recorded at 23°C (black) and 31°C (red) were normalized to their peak value. Single-exponential fits to the decay phase of averaged EPSCs showed that τ_decay_ were 0.47 ms at 24°C (black dash line) and 0.26 ms at 31°C (red dash line), respectively.

**Table 2.** Biophysical properties of afferent fiber spontaneous EPSCs and EPSPs

| **EPSC (n = 18)** | **23.4 ± 0.1°C** | **31.1 ± 0.1°C** | ***p* value** | **Q<sub>10</sub>** |
| --- | --- | --- | --- | --- |
| **Amplitude (pA)** | 80.5 ± 6.8 | 77.7 ± 7.4 | 0.61 | 1.1 ± 0.2 |
| **Frequency (Hz)** | 78.2 ± 9.5 | 115.2 ± 9.7 | 0.0097 | 3.2 ± 1.1 |
| **Rise time (μs)** | 223 ± 13 | 150 ± 6 | <0.0001 | 1.7 ± 0.1 |
| **τ<sub>decay</sub> (μs)** | 617 ± 36 | 400 ± 24 | <0.0001 | 1.8 ± 0.1 |
| **Charge transfer (fC)** | -71.3 ± 5.2 | -47.9 ± 3.4 | <0.0001 |  |
| **EPSP (n = 19)** | **23.4 ± 0.1°C** | **31.5 ± 0.1°C** | ***p* value** | **Q<sub>10</sub>** |
| **Amplitude (mV)** | 3.9 ± 0.3 | 3.8 ± 0.4 | 0.8 | 1.1 ± 0.2 |
| **Frequency (Hz)** | 40.4 ± 6.1 | 56.9 ± 7.0 | 0.003 | 2.1 ± 0.5 |
| **τ<sub>decay</sub> (ms)** | 3.50 ± 0.69 | 1.67 ± 0.38 | <0.0001 | 2.7 ± 0.4 |

Values are means ± SEM

Among all the afferent fibers recorded, 5 out of 18 (28%) exhibited double peaks in the EPSCs amplitude distribution at both room and high temperatures. Fig. 2B1 shows spontaneous EPSCs recorded from an afferent fiber that had a double peak distribution (Fig. 2B2). Double or triple peak EPSC amplitude distributions have been observed previously in afferent fibers of rats (Grant et al., 2010) and turtle (Schnee et al., 2013). We divided EPSCs into two groups according to their amplitude: small EPSCs with amplitude <150 pA and large EPSCs with amplitude >150 pA. The frequency distributions of EPSC amplitude in each group were fit well with a Gaussian function (small events: R^2^ = 0.94 at 23°C, R^2^ = 0.93 at 31°C; large events: R^2^ = 0.93 at 23°C, R^2^ = 0.97 at 31°C). Under 23°C, the 1^st^ and the 2^nd^ peak of the Gaussian fits were 78 pA and 292 pA, respectively. Under 31°C, the 1^st^ and the 2^nd^ peak were 89 pA and 279 pA, respectively. Under 23°C, there were 65% of EPSCs larger than 150 pA. The group of large EPSCs increased to 74% under 31°C.

Spontaneous EPSPs were also recorded from afferent fibers using whole cell current clamp with 0 pA holding current. The response to temperature changes of the EPSP amplitudes was variable: Figure 3A shows one example with decreased EPSP amplitude at high temperature and Figure 3B shows another example with increased EPSP amplitude at high temperatures. The EPSP amplitude distribution is shown in Fig. 3A2, where 1250 and 1817 EPSPs events were counted from an afferent fiber under 23°C and 31°C, respectively. The amplitude distributions of EPSPs were fit well with a Gaussian function (Fig. 3A2: R^2^ = 0.89 at 23°C; R^2^ = 0.97 at 31°C). The peaks of the Gaussian fit are 5.7 mV at 23°C and 2.7 mV at 31°C. The single-exponential fit to the decay phase of averaged EPSPs (Fig. 3A3) showed that τ_decay_ was 2.7 ± 0.4 ms at room temperature (n = 19, Table 2). This is faster than that found in afferent fibers of immature rat (Yi et al, 2010). Spontaneous EPSPs shown in Fig. 3B1 were recorded from the same afferent fiber with double peak EPSC amplitude distribution as shown in Fig. 2B2. During 30 second long recordings, 1054 and 1581 EPSPs were recorded from another afferent fiber under room and high temperature, respectively (Fig. 3B2). The frequency distribution of EPSP amplitudes shown in Fig. 3B2 also had double peaks at high temperature, but not under room temperature. Under 23°C, the peak of the Gaussian fit was 2.7 mV (R^2^ = 0.87). Under 31°C, the 1^st^ and the 2^nd^ peak were 3.4 mV (R^2^ = 0.85) and 7.3 mV (R^2^ = 0.81), respectively.

**Fig. 3.**
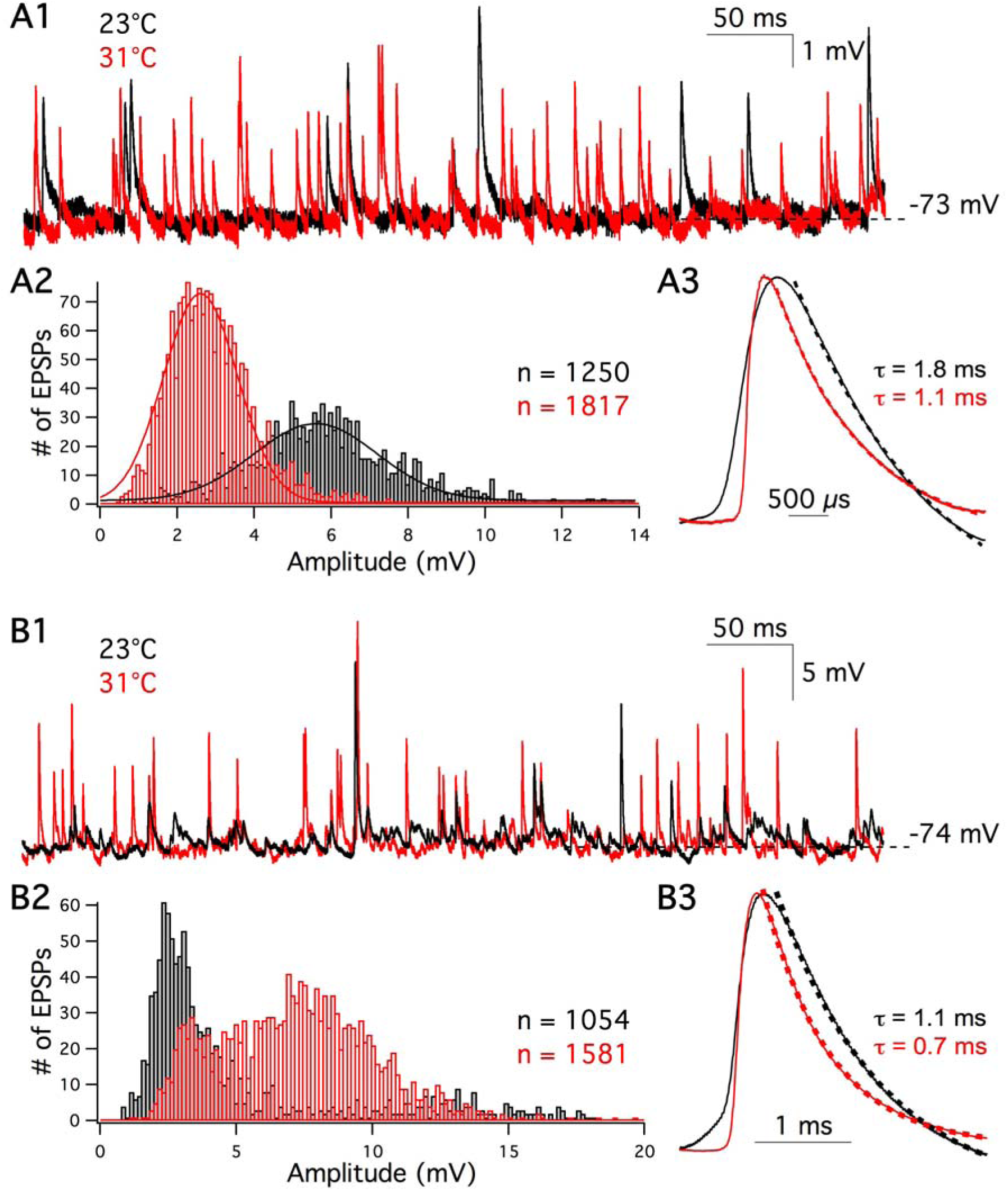
EPSPs are faster and their frequency increases at high temperature. (A1) Spontaneous EPSPs were recorded from an afferent fiber at 23°C (black) and 31°C (red). The membrane current of fiber was clamped at 0 pA. (A2) A total of 1250 and 1817 EPSPs were recorded at 23°C (black) and 31°C (red), respectively. The EPSP amplitude distributions were fit with a Gaussian function. (A3) Average EPSPs recorded at 23°C (black solid line) and 31°C (red solid line) were normalized to their peak values. The dashed lines are single-exponential fits to the decay phase of the average EPSPs. In this example, τ_decay_ was 1.8 ms at 23°C (black dash line) and 1.1 ms at 31°C (red dash line), respectively. (B1) Spontaneous EPSPs were recorded from the same afferent fiber as shown in Fig. 2 (B1 to B3) at 23°C (black) and 31°C (red). (B2) A total of 1054 and 1581 EPSPs were recorded under room and high temperature. (B3) Average of EPSPs recorded at 23°C (black) and 31°C (red) were normalized to their peak values. Single-exponential fits to the decay phase of averaged EPSPs (dash lines) revealed that τ_decay_ was 1.1 ms at 23°C and 0.7 ms at 31°C, respectively.

**Fig. 4.**
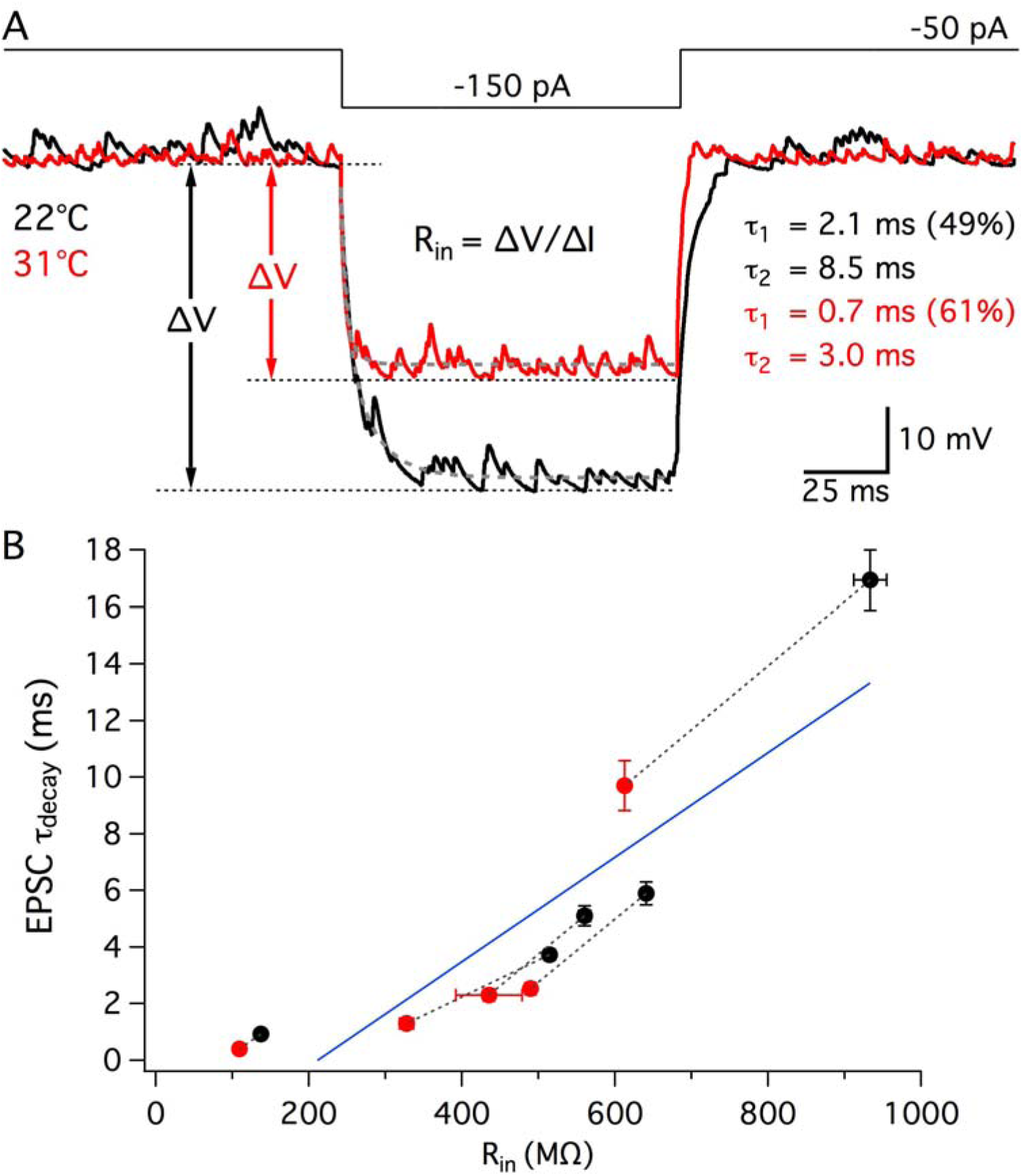
Afferent fiber membrane input resistance and EPSP decay kinetics. (A) In current clamp, afferent fibers were hyperpolarized by injection of negative currents from −50 pA to −150 pA for 100 ms. Averages of membrane potential responses from one afferent fiber under 22°C and 31°C are shown in black and red, respectively. Red and black arrows indicate the absolute change in membrane potentials at 22°C and 31°C, respectively. Input resistance of an afferent fiber was calculated following Ohm’s law, R_in_ = ΔV/ΔI. The decay phase of the membrane potential was fit by a double exponential function (gray dashed lines). (B) τ_decay_ of spontaneous EPSPs was plotted against R_in_ of the respective fibers. Each pair of data points recorded from the same afferent fiber is connected with a dashed line: room (black) and high (red) temperature (n = 5). A linear regression fit is shown in blue and indicates a positive correlation between EPSP τ_decay_ and R_in_ (Slope = 0.018 ± 0.003, R^2^ = 0.79, *p* = 0.0006).

**Fig. 5.**
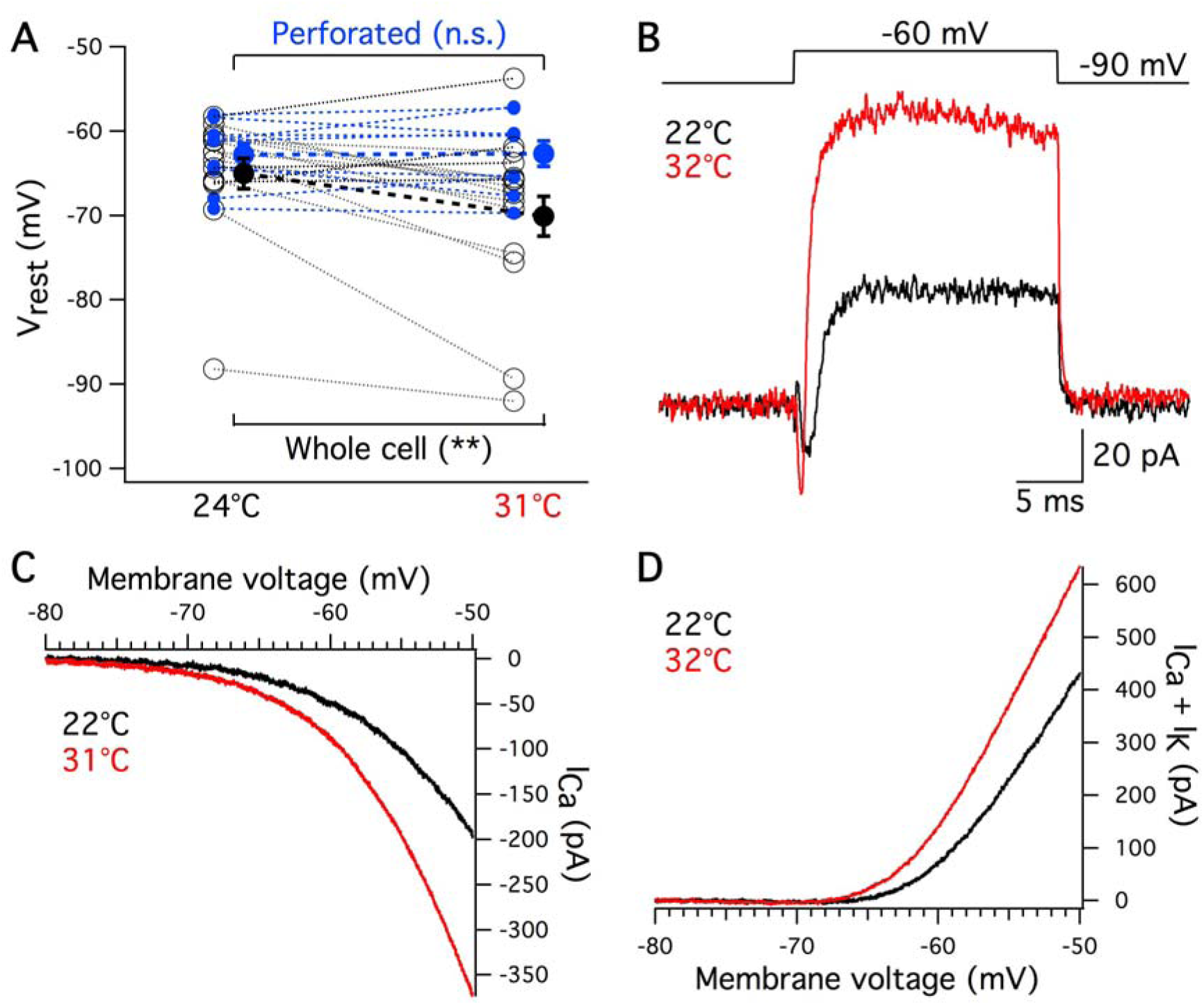
Hair cell resting membrane potential and ionic currents at high temperature. (A) A K^+^-based internal pipette solution and current-clamp recordings with zero current injection were used to measure the V_rest_ of hair cells. Whole-cell recordings showed that V_rest_ hyperpolarized at high temperature (black circles, *p* = 0.006, n = 15). Gramicidin-mediated perforated patch recordings showed that V_rest_ remained the same at high temperature (blue dots, *p* = 0.88, n = 9). (B) Whole-cell voltage-clamp recordings of hair cells with K^+^-based internal solution showed current changes in response of a step depolarization from −90 mV to −60 mV. The traces were averaged from five hair cells recorded at 22°C (black) and 32°C (red). Note the faster kinetics of the currents at 32°C. (C) In order to reveal Ca^2+^ current, hair cells were patched with a Cs^+^-TEA-based internal solution. Hair cells were depolarized with a ramp depolarization from −90 mV to −50 mV for 400 ms at 22°C (black) and 31°C (red). The traces were averaged from five hair cells. (D) Voltage-dependent K^+^ and Ca^2+^currents were triggered by the same ramp stimulus with K^+^-based internal solution. The traces were averaged from fourteen hair cells recorded at 22°C (black) and 32°C (red). Note how the larger outward K^+^ currents dominate the hair cell I-V curve.

**Fig. 6.**
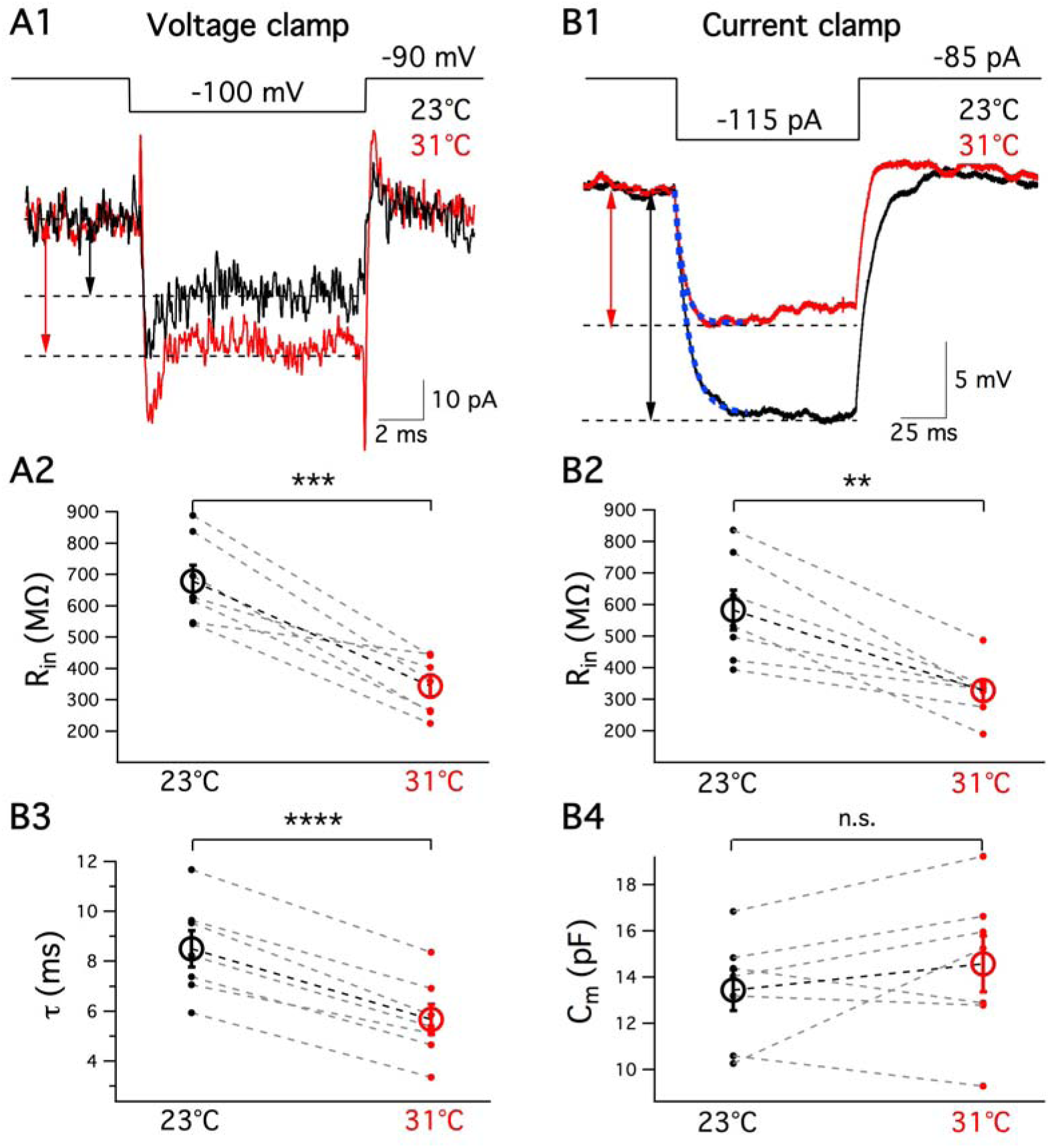
Temperature effects on hair cell passive membrane properties. (A1) Inward currents were triggered by a small hyperpolarization from −90 mV to −100 mV under 23°C (black) and 31°C (red). The size of the current was the difference between baseline and the plateau inward current (dashed lines). (B1) Hyperpolarized membrane potential in response to a −30 pA current injection under current clamp were recorded at 23°C (black) and 31°C (red). The change in membrane potential was calculated by subtracting baseline from plateau (dashed lines). (A2) Using voltage clamp measurements and R_in_ = ΔV/ΔI, the membrane input resistance (R_in_) was lower at high temperature (*p* = 0.0006, n = 7). (B2) Current clamp measurements also showed that R_in_ decreased at high temperature (*p* = 0.0031, n = 7). (B3) Membrane time constant (τ) was estimated by fitting a single exponential to a 40-ms time window of the voltage response following the initial current injection (superimposed blue dash lines in B1). τ was faster at high temperature (p < 0.0001, n = 7). (B4) C_m_ calculated using C_m_ = τ/R_in_ did not show significant change at high temperature (*p* = 0.2506, n = 7). In (A2), (B2) to (B4), individual values, mean and SEM are shown in dots, open circles and bars, respectively.

**Fig. 7.**
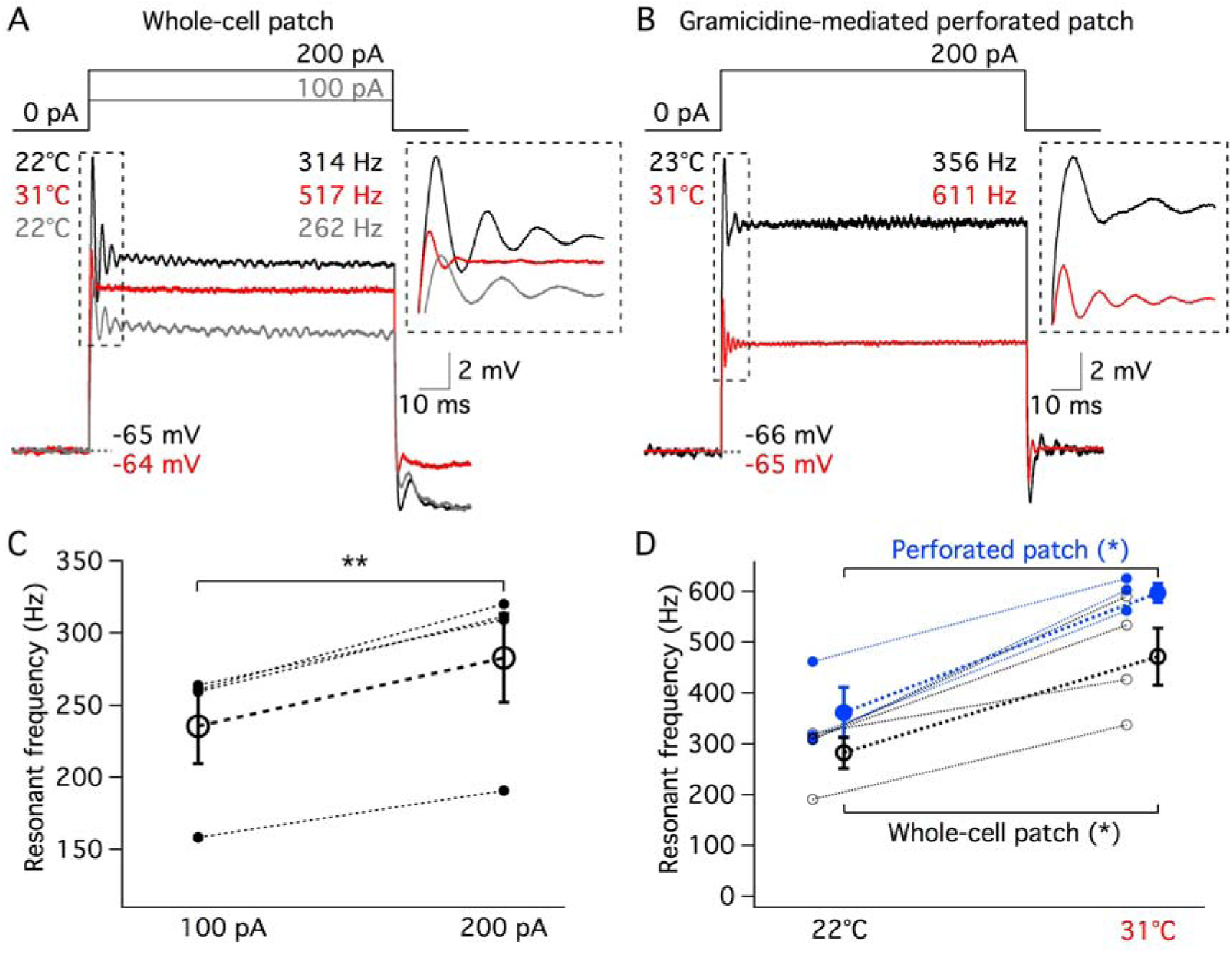
Hair cell resonant frequency increases at high temperature. (A) Electrical resonance was elicited by stimulating hair cells with a current step in whole-cell patch. Black and red traces were evoked by a +200 pA current step at room and high temperature, respectively. Gray trace was evoked by a +100 pA current step at room temperature. Each of these three traces is the average response from three hair cells. Resonant frequencies were calculated by fitting the first 20-ms membrane voltage changes with a sine wave function. The resonant frequencies of black, red and gray traces were 314 Hz, 517 Hz and 262 Hz, respectively. (B) Electrical resonance of hair cells was elicited by a +200 pA current step in gramicidine-mediated perforated patch. Black and red traces were averaged of 3 cells recorded at room and high temperature, respectively. The resonant frequencies of black and red traces were 356 Hz and 611 Hz, respectively. (C) Resonant frequencies of hair cells evoked by +100 pA current steps were lower than those evoked by +200 pA current steps (n = 4, *p* = 0.004). (D) Results of both whole-cell (black, n = 4, *p* = 0.017) and perforated (blue, n = 3, *p* = 0.025) patch recordings showed that resonant frequency of hair cells increased significantly at high temperature. There was no statistical difference between resonant frequencies measured by whole-cell patch or perforated patch at either room (*p* = 0.80, unpaired *t* test) or high (*p* = 0.56, unpaired *t* test) temperature.

The V_rest_ of the afferent fiber was slightly depolarized from −74.0 ± 0.7 mV at 23°C to −71.2 ± 0.8 mV at 32°C (*p* = 0.0012, n = 19). Consistent with EPSC frequency, the frequency of the EPSPs increased at high temperature (Table 2). Averaging the entire data set revealed no statistical difference in the amplitude of EPSPs between room and high temperature (3.9 ± 0.3 mV vs 3.8 ± 0.4 mV, respectively; n = 19, *p* = 0.8, Table 2). However, 58% (11/19) of the fibers showed a reduced EPSP amplitude at high temperature.

### Afferent fiber membrane input resistance at high temperature

The τ_decay_ of EPSPs decreased at high temperature in all of the fibers recorded (Fig. 3B3; Table 2), which may due to lower membrane input resistance (R_in_) at high temperature. We thus estimated R_in_ of the afferent fibers using current clamp: R_in_ = ΔV/ΔI, where ΔV was the change in membrane potential induced by a 100 pA current injection (Fig. 4A). The R_in_ of afferent fibers decreased from 557 ± 128 M at 22.0 ± 0.1°C to 395 ± 85 MΩ at 32.0 ± 0.2°C (n = 5, *p* = 0.027). In this group of fibers, τ_decay_ of spontaneous EPSPs were 6.5 ± 2.7 ms and 3.3 ± 1.7 ms at room and high temperature, respectively (n = 5, *p* = 0.041). The linear regression (Slope = 0.018 ± 0.003, R^2^ = 0.79, *p* = 0.0006) indicated that EPSP τ_decay_ is positively correlated with afferent fiber R_in_ (Fig. 4B, blue line). An afferent fiber is composed of two compartments: a calyx-type ending that contacted the hair cell and a thin fiber cable. We thus fit the decay phase of membrane potential change to a double exponential function (Fig. 4A gray dashed lines that superimposed over V_m_ curves):

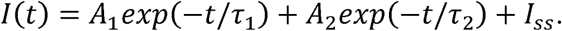

We can then calculate a weighted mean time constant *τ_mean_* using:

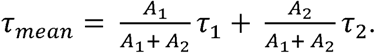

Consistent with R_in_, *τ_mean_* was shorter at high temperature (4.8 ± 1.5 ms) than that at room temperature (7.8 ± 1.5 ms, n = 5, *p* = 0.0014). However, EPSP amplitude did not change significantly with temperature for this data set (6.5 ± 0.7 mV vs 9.3 ± 2.2 mV at 22.1 ± 0.2°C and 32.0 ± 0.1°C, respectively; n = 5, *p* = 0.3). The linear regression of EPSP amplitude and R_in_ also did not show a significant difference (slope = −0.0016 ± 0.005, R^2^ = 0.013, *p* = 0.76). In conclusion, the R_in_ of afferent fibers exhibit heterogeneity from 100 MΩ to 1000 MΩ which could contribute to the *in vivo* variation in spontaneous spike rates and afferent fiber thresholds to sound stimulation (Stiebler and Narins, 1990).

Changes in R_in_ with temperature are also important because of membrane potential noise considerations (Fatt and Katz, 1952; Johnson, 1987). The mean square membrane voltage is given by the Nyquist theorem:

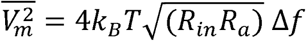

here *k_B_* is Boltzmann constant, T the temperature in degrees Kelvin (T= 273 + temperature in Celsius), *R_in_* the membrane input resistance, *R_a_* the axial cable resistance and Δ*f* the noise frequency bandwidth. The frequency bandwidth is inversely proportional to the membrane time constant, which determines the cell cut-off frequency, and will thus increase at higher temperatures. The decrease in R_in_ with increasing temperature is thus important because it counteracts the increase in noise bandwidth with temperature. This helps to keep the membrane potential in a stable sub-threshold level.

### Resting membrane potential of hair cells

The increase in spontaneous EPSC frequency at high temperature suggests more spontaneous vesicle release from hair cells, which may be due to a depolarization of V_rest_ at high temperature. We thus measured hair cell V_rest_ under whole-cell current clamp with a holding current at 0 pA. Unexpectedly, whole-cell recordings revealed that V_rest_ hyperpolarized from −65.0 ± 1.8 mV at 23.2 ± 0.3°C to −70.1 ± 2.3 mV at 31.2 ± 0.3°C (*p* = 0.006, n = 15, Fig. 5A, black circles). To prevent the dialysis of the hair cell constituents through the patch pipette, we also performed perforated-patch recordings using gramicidin (40-50 μg/ml). The perforated patch clamp showed that V_rest_ did not change at high temperatures (V_rest_ = −62.8 ± 1.3 mV at 23.4 ± 0.5°C and V_rest_ = −62.7 ± 1.5 mV at 31.6 ± 0.2°C, *p* = 0.88, n = 9; Fig. 5A, blue dots).

### Hair cell potassium and calcium currents at high temperature

Under voltage clamp and with a K^+^-based internal pipette solution, we determined the voltage-dependent currents that are activated by a step-depolarization from −90 mV to −60 mV. A rapidly activating inward current was followed by a rapid and large outward current (Fig. 5B). The inward current was a Ca^2+^ current that is activated more quickly at higher temperatures in frog hair cells (Hudspeth and Lewis, 1988). Using Cs^+^-TEA-based or K^+^-based internal solutions, we recorded the I-V curves of Ca^2+^ currents (Fig. 5C) and the overall K^+^ + Ca^2+^ currents (Fig. 5D), respectively. Both types of currents increased in amplitude at high temperature. Importantly, note that the slope of voltage-dependent Ca^2+^ current was greater at high temperature making Ca^2+^ influx more sensitive to small membrane potential changes. The increase in whole cell outward K^+^ + Ca^2+^ current at high temperature explains the decrease in V_rest_ with temperature (Fig. 5A, black circles). In perforated patch recordings a lack of change in V_rest_ could be due to a simultaneous and more compensatory increase in inward Ca^2+^ current and outward K^+^ current as temperature increases.

### Hair cell membrane input resistance decreases at high temperature

At high temperature an increase in Ca^2+^ and K^+^ currents around V_rest_ should result in a decrease in membrane input resistance (R_in_). To check this we performed whole-cell voltage and current clamp recordings from hair cells. In voltage clamp, hyperpolarizing hair cells from −90 mV to −95 mV or −100 mV for 10 ms elicited small inward currents (Fig. 6A1). R_in_ was calculated from Ohm’s law (R_in_ = ΔV/ΔI). R_in_ of hair cells decreased at high temperature (659.7 ± 54.3 MΩ at 23.2 ± 0.2°C and 343.7 ± 34.7 MΩ at 31.4 ± 0.2°C, *p* = 0.0006, n = 7, Fig. 6 A2). Under current clamp, injection of −20 pA or −30 pA hyperpolarized the cells (Fig. 6 B1). Current clamp also showed a decrease in R_in_ at high temperature (640.3 ± 56.4 MΩ at 23.2 ± 0.2°C and 389.3 ± 28.2 MΩ at 31.2 ± 0.2°C, *p* = 0.0023, n = 7, Fig. 6 B2). Changes in current or membrane potential were determined by calculating the differences between baseline and plateau (dashed lines in Fig. 6A1 and 5B1). There was no statistically significant difference in R_in_ measured with voltage clamp or current clamp at either room (*p* = 0.2598) or high (*p* = 0.7016) temperature (n = 7).

Some hair cells tend to be leakier towards the end of recordings. The leak current contributes to cell conductance and affects R_in_ (Ceballos et al., 2017). To exclude the influence of the recording order on R_in_, we measured R_in_ in another group of hair cells. In eight cells, we first recorded at room temperature and then heated the solution to high temperature. R_in_ decreased from 578.8 ± 49.8 MΩ at 24.0 ± 0.2°C to 315.8 ± 33.3 MΩ at 31.5 ± 0.2°C (p = 0.0001, n = 8). In another three cells, we recorded first at high temperature and then at room temperature. R_in_ was also lower at high temperature (600.2 ± 38.2 MΩ at 24.1 ± 0.1°C and 277.6 ± 34.0 MΩ at 31.9 ± 0.1°C, *p* = 0.0146, n = 3). Thus, the order of recording did not affect R_in_ measured at either room (*p* = 0.81) or high (*p* = 0.53) temperature. This suggests that the decrease in R_in_ at high temperature was not due to hair cells becoming leakier towards the end of the recordings, but due to an increase in cell membrane conductance at high temperature (see Fig. 5B).

Using current clamp data, we also calculated the membrane time constant (τ). We fitted a single exponential to a 40-ms time window of the voltage response following the initial current injection (Fig. 6B1, superimposed blue dashed line). The time constant was 8.50 ± 0.73 ms at 23.2 ± 0.2°C and was 5.67 ± 0.61 ms at 31.2 ± 0.2°C (*p* < 0.0001, n = 7, Fig. 6B3). We next determined the membrane capacitance (C_m_) using the equation C_m_ = τ/R_in_. The average value of C_m_ remained the same regardless of temperature changes (13.5 ± 0.9 pF at 23.2 ± 0.2°C and 14.6 ± 1.2 pF at 31.2 ± 0.2°C, *p* = 0.2506, n = 7; Fig. 6B4). This value of C_m_ is similar to previous whole-cell recordings in voltage clamp of C_m_ in bullfrog hair cells (Li et al., 2009).

### Electrical resonant frequency increases at high temperature

Unlike mechanical tuning in the mammalian cochlea, amphibian papilla hair cells in frogs are electrically tuned (Smotherman and Narins, 2000). Electrical resonance of amphibian papilla hair cells matches the characteristic (best) frequencies of the afferent fibers (Smotherman and Narins, 1999). The electrical resonance is a result of the interplay of voltage-gated ion channels and membrane capacitance (Crawford and Fettiplace, 1981; Art and Fettiplace, 1987; Hudspeth and Lewis, 1988). Since the conductance of voltage-gated calcium and potassium channels are temperature dependent as shown in Fig. 5, we expect electrical resonance to be temperature sensitive. To elicit electrical ringing around resting membrane potentials, hair cells were injected a current step from zero current under whole-cell (Fig. 7A) or perforated patch clamp (Fig. 7B). Resonant frequency was estimated by fitting the first 20-ms of the damped oscillation in membrane voltage with a sine wave function (Li et al., 2014). In a previous study (Smotherman and Narins, 1999), it was shown that resonant frequency increases as larger depolarizing current steps are applied to the cell. Consistently, depolarizing hair cells with +100 pA current steps elicited a resonant frequency of 236 ± 26 Hz, while +200 pA current steps elicited a higher resonant frequency of 283 ± 31 Hz at room temperature (*p* = 0.0038, n = 4; Fig. 7A and 7C). To compare the resonant frequency measured with whole cell recordings to perforated patch recordings, we chose the larger +200 pA current steps. The resonant frequency increased from 283 ± 31 Hz at 21.8 ± 0.4°C to 472 ± 56 Hz at 31.4 ± 0.6°C (*p* = 0.017, n = 4, Fig. 7D black circle). Similarly, perforated patch recordings revealed that resonant frequency increased from 362 ± 50 Hz at 24.8 ± 2.1°C to 597 ± 19 Hz at 31.5 ± 0.3°C (*p* = 0.025, n = 3, Fig. 7D blue dots). There was no statistical difference in resonant frequency measured by whole-cell recording and perforated patch at either room (*p* = 0.22, unpaired *t* test) or high (*p* = 0.13, unpaired *t* test) temperatures.

Hair cell resonant frequency also increases when resting membrane potential is depolarized (Hudspeth and Lewis, 1988; Smotherman and Narins, 1999). However, the resting membrane potential did not change for the whole cell recordings of Figure 7A (−65.2 ± 0.5 mV at room temperature vs. −63.8 ± 0.8 mV at high temperature, *p* = 0.23, n = 4). Similarly, resting membrane potential did not change with temperature for the perforated patch data (−66.0 ± 2.7 mV at room temperature vs. −65.2 ± 2.6 mV at high temperature, *p* = 0.51, n = 3). Therefore, the increases of resonant frequency at high temperature are not due to changes in resting membrane potential. Instead, they are caused by faster activation and larger conductance of Ca^2+^ and K^+^ channels upon depolarization at high temperature (Hudspeth and Lewis, 1988; Smotherman and Narins, 1999). The resonant frequency measured at room temperature matches the results of *in vitro* whole cell recordings of hair cells from amphibian papilla using a ZAP protocol (Frolov and Li, 2017). The Q_10_ of resonant frequency of amphibian papilla hair cells (1.7 ± 0.1, n = 4 for whole cell; 1.9 ± 0.08, n = 3 for perforated patch) are similar to that recorded from frog saccular hair cells (Smotherman and Narins, 1998). This temperature-dependent shift in hair cell resonant frequency may be responsible for the temperature-induced shifts of best frequency observed at *in vivo* amphibian papilla auditory nerve fiber recordings (Stiebler and Narins, 1990).

### Release efficiency increases at high temperature

A lower R_in_ at high temperature results in smaller membrane potentials changes, which will decrease hair cell sensitivity to low-level sounds. To explore whether hair cells compensate for this putative loss in sensitivity, we studied the temperature dependence of vesicle exocytosis from hair cells using time-resolved membrane capacitance measurements (Lindau and Neher, 1988; Gillis, 2000). Changes in C_m_ (ΔC_m_) were evoked by depolarizing voltage steps. We first employed 20-ms depolarizing steps from a holding potential of −90 mV to −30 mV to evoke fast release that depletes the readily releasable pool (RRP) consisting of vesicles docked near the clustered Ca^2+^ channels (Graydon et al., 2011; Fig. 8A1). The depolarization triggered a larger Ca^2+^ current (Fig. 8A1) under high temperature than under room temperature (*p* <0.0001, n = 7): Ca^2+^ charge transfer (Q_Ca_) was 11.9 ± 1.3 pC at 23.3 ± 0.2°C and that was 15.9 ± 1.7 pC at 32.0 ± 0.3°C. The Q_10_ of 20-ms pulse evoked Q_Ca_ was 1.5 ± 0.05 (n = 7). Average resting membrane capacitance measured at high temperature (16.9 ± 0.7 pF) was similar to that measured at room temperature (15.7 ± 0.6 pF, *p* = 0.0841, n = 7). The ΔC_m_ increased from 27.2 ± 4.7 fF at room temperature to 51.6 ± 6.0 fF at high temperature (*p* = 0.0006, n = 7). The Q_10_ of 20-ms pulse evoked ΔC_m_

Here Q_10_ is the ratio of the reaction rates for T_L_ = 296°K (23°C) and for T_H_ = T_L_ +10°K = 306°K:

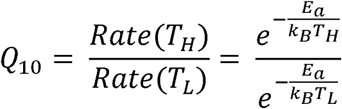

where *k_B_* ≈ 8.3 J/mol°K (Boltzmann constant) and E_a_ is the reaction activation energy barrier. From this equation we obtain *E_a_* = (*lnQ*_10_)·75.2 kJ/mol. The E_a_ for evoked exocytosis was thus 65.8 ± 10.0 kJ/mol ≈ 26.8 k_B_T (n = 7), which was calculated from the Q_10_ of 20-ms depolarizing pulses. This is within the E_a_ estimated for vesicle exocytosis mediated by SNAREpin and synaptotagmin molecules (20 to 140 k_B_T; Mostafavi et al., 2017; Schotten et al., 2015; Zhang and Jackson, 2008), although exocytosis in hair cells is mediated by otoferlin (Michalski et al., 2017).

**Fig. 8.**
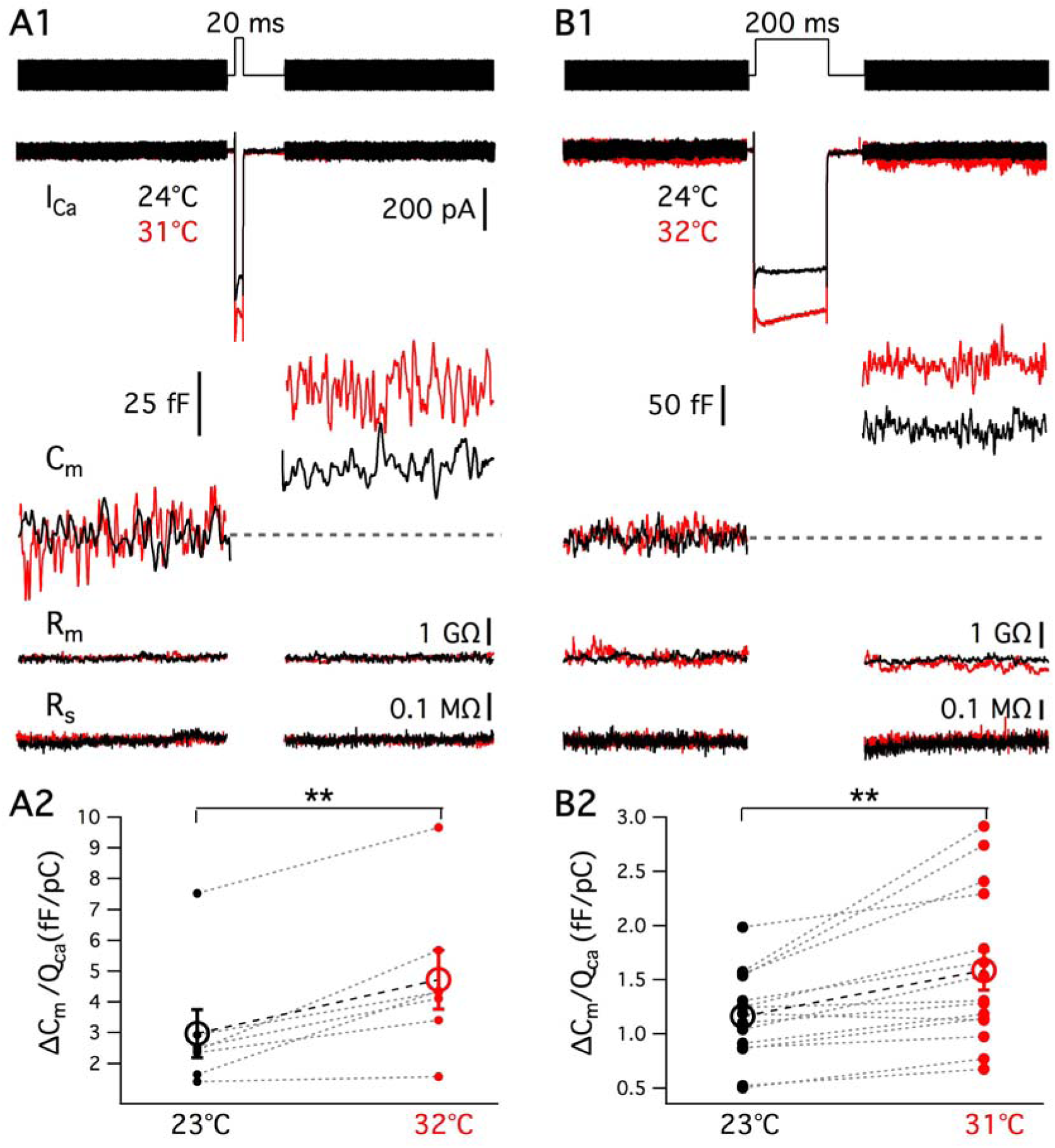
Hair cell exocytosis efficiency is enhanced at high temperature. (A1) and (B1) show the Ca^2+^ current (I_Ca_), membrane capacitance (C_m_), membrane resistance (R_m_) and series resistance (R_s_) traces obtained from a hair cell. A ΔC_m_ change was triggered by 20 ms (A1) and 200 ms (B1) depolarizing steps from −90 mV to −30 mV at room (black) and high (red) temperature. For the 20-ms depolarization: I_hold_ = −60 pA and −64 pA; C_m(resting)_ = 14.5 pF and 16.0 pF; R_m_ = 1.3 GΩ and 1.2 GΩ; R_s_ = 8.9 MΩ and 8.2 MΩ, at 24°C and 31°C, respectively. For recordings of 200-ms depolarization: I_hold_ = −69 pA and −43 pA; C_m(resting)_ = 11.3 pF and 12.8 pF; R_m_ = 1.2 GΩ and 1.5 GΩ; R_s_ = 14.0 M and 11.3 MΩ at 24°C and 32°C, respectively. Exocytosis efficiency, calculated as the ratio of ΔC_m_/*Q_Ca_*^2+^, is shown in (A2) and (B2) for 20-ms (*p* = 0.0043, n = 7) and 200-ms (*p* = 0.0013 n = 15) depolarizations, respectively. The exocytosis efficiency is nearly 3-fold greater for the short 20 ms pulse than for the long 200 ms pulse.

**Fig. 9.**
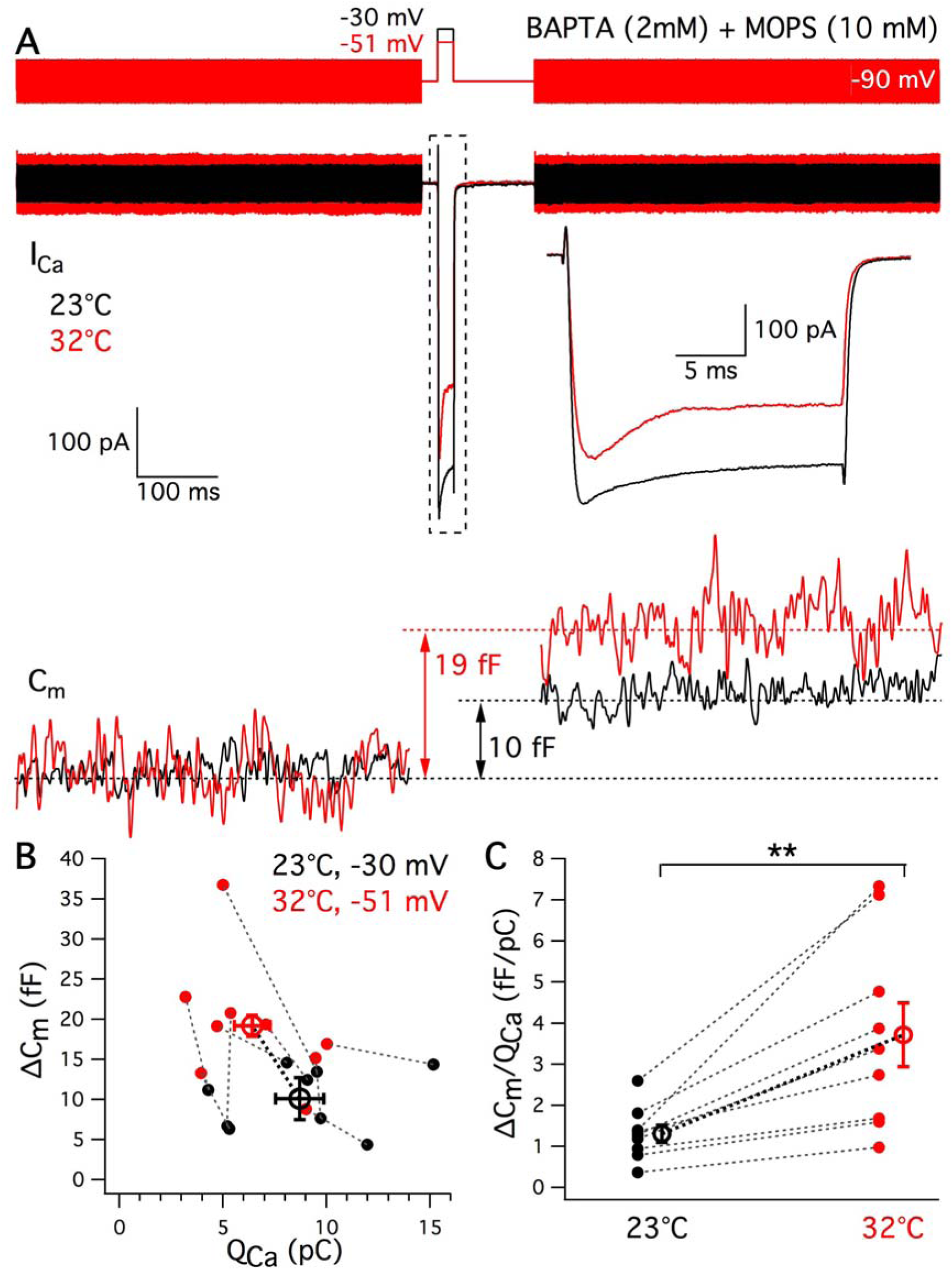
Exocytosis increases at high temperature even when Ca^2+^ influx is reduced. (A) The depolarization potential at high temperature was adjusted from −45 to −55 mV to elicit the same or less amount of Ca^2+^ influx to that at room temperature. In order to exclude temperature effects on pH or Ca^2+^ buffering, hair cells were patched with internal solution containing pH-insensitive Ca^2+^ buffer BAPTA (2 mM) and a different pH buffer MOPS (10 mM). Averages of Ca^2+^ currents (I_Ca_) from nine hair cells triggered by depolarization to −30 mV at 23°C (black) and to −51 ± 2 mV at 32°C (red). The I_Ca_ was expanded in time scale from the one in a dash line box. The membrane capacitance change (ΔC_m_) was 10 fF at 23°C (black) and 19 fF at 32°C (red). (B) ΔC_m_ versus Q_Ca_ from individual hair cells depolarized to −30 mV at 23°C (black) and to −51 mV at 32°C (red). Data points connected by a dashed line were recorded from the same hair cell. (C) Exocytosis efficiency (ΔC_m_/*Q_ca_*^2+^) increased from 23°C (black) to 32°C (red; *p* = 0.0042, n = 9).

Depolarizations from −90 mV to −30 mV for 200 ms (Fig. 8B1) not only depletes the RRP but also evokes sustained release of vesicles replenishing to the RRP (Graydon et al., 2011). Q_Ca_ evoked by the 200-ms depolarizing pulse was 99.7 ± 8.0 pC at 23.1 ± 0.2°C and that was 126.3 ± 9.7 pC at 31.3 ± 0.2°C (p <0.0001, n = 15). ΔC_m_ evoked by 200-ms depolarizing steps increased from 109.8 ± 9.1 fF at room temperature to 192.1 ± 21.2 fF at high temperature (*p* < 0.0001, n = 15). The Q_10_ of Q_Ca_ and ΔC_m_ evoked by a 200-ms pulse were 1.3 ± 0.04 and 2.0 ± 0.1 (n = 15), respectively. The Q_10_ of exocytosis from bullfrog hair cells is close to that found in mouse inner hair cells using capacitance measurements (Q_10_ = 2.1; Nouvian, 2007) and in rat hippocampal neurons using FM1-43 destaining methods (Q_10_ = 2.5; Yang et al., 2005).

Plotting ΔC_m_ versus the corresponding *Q_ca_*^2+^, we obtained the efficiency of exocytosis as the ratio of ΔC_m_/*Q_ca_*^2+^, which for 20-ms pulses increased from 3.0 ± 0.8 fF·pC^−1^ at room temperature to 4.7 ± 1.0 fF·pC^−1^ at high temperature (*p* = 0.0043, n = 7; Q_10_ = 1.9 ± 0.3; Fig. 8 A2). The ratio of ΔC_m_/*Q_ca_*^2+^ for responses to 200-ms pulses increased from 1.2 ± 0.1 fF pC^−1^ at room temperature to 1.6 ± 0.2 fF pC^−1^ at high temperature (*p* = 0.0013, n = 15; Fig. 8 B2). Recordings from mouse IHC also find that higher temperatures enhance Ca^2+^ influx and increase ΔC_m_/*Q_ca_*^2+^ elicited by 20-ms depolarizing pulses, but not for sustained exocytosis (Nouvian, 2007). By contrast, our results with bullfrog hair cells indicate that the efficiency of both fast and sustained exocytosis increased under high temperature.

To further demonstrate that the increase in exocytosis is facilitated beyond a mere increase of Ca^2+^ influx under high temperature, we reduced Ca^2+^ influx using smaller depolarizing potentials at high temperature and measured ΔC_m_ jumps. We also employed the pH-insensitive Ca^2+^ buffer BAPTA (Tsien, 1980) and a different pH buffer MOPS, which has similar temperature dependence with HEPES (Ellis and Morrison, 1982). A depolarization from −90 mV to potential that varied −46 to −56 mV (average: −50.6 ± 1.8 mV, n = 9) for 20 ms at 32°C elicited similar or less Ca^2+^ influx (6.4 ± 0.8 pC) to that triggered by 20-ms depolarization to −30 mV under 23°C (8.7 ± 1.2 pC, *p* = 0.0059, n = 9; Fig. 9A). However, even when Ca^2+^ influx was reduced hair cells were still able to release more vesicles upon depolarization at high temperature (Fig. 9A). ΔC_m_ increased from 10.3 ± 1.3 fF at room temperature to 19.2 ± 2.6 fF at high temperature (*p* = 0.0028, n = 9, Fig. 9B). Consistent with previous results using internal EGTA and HEPES, exocytosis efficiency increased from 1.3 ± 0.2 at room temperature to 3.7 ± 0.8 at high temperature (*p* = 0.0042, n = 9; Fig. 9C). The results indicate that high temperature facilitates synaptic vesicle exocytosis even when Ca^2+^ influx is reduced. Moreover, the effect is not due to temperature dependent changes in cellular pH-dependent Ca^2+^ buffering.

### Readily releasable pool (RRP) size is larger at high temperature

To compare the maximum size of hair cell RRP under room and high temperature, we employed a dual-pulse protocol (Gillis et al., 1996). Two pulses of 100 ms duration are given with a 25-ms interval (Fig. 10A). The short 25-ms interval results in depression in the second release. Ca^2+^ current can also decline during the dual pulse due to inactivation. To avoid differences in Q_Ca_ during the first and second depolarization, the potentials of the first pulses are adjusted (Fig. 10A). The average of the first depolarizing potentials were −41.1 ± 0.7 mV (n = 11). The intervals between pulse pairs was 60-90 seconds to allow for full recovery. Under 23°C, the Ca^2+^ influx during the first and second depolarizing pulses were 31.8 ± 6.8 pC and 31.7 ± 6.8 pC, respectively (n = 4, *p* = 0.93, Fig. 10 A and C). Under 31°C, the Ca^2+^ influx during the first and second depolarizing pulses were 62.8 ± 3.8 pC and 63.3 ± 3.1 pC, respectively (n = 7, *p* = 0.69, Fig. 10 A and C). Consistently, both the first (*p* = 0.0019) and the second (*p* = 0.0009, Fig. 10C) pulses triggered more Ca^2+^ influx under high temperature. ΔC_1_ and ΔC_2_ were capacitance jumps triggered by the first and second depolarization. The maximum size of the RRP (B_max_) was calculated according to:

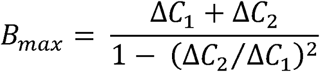

**Fig. 10.**
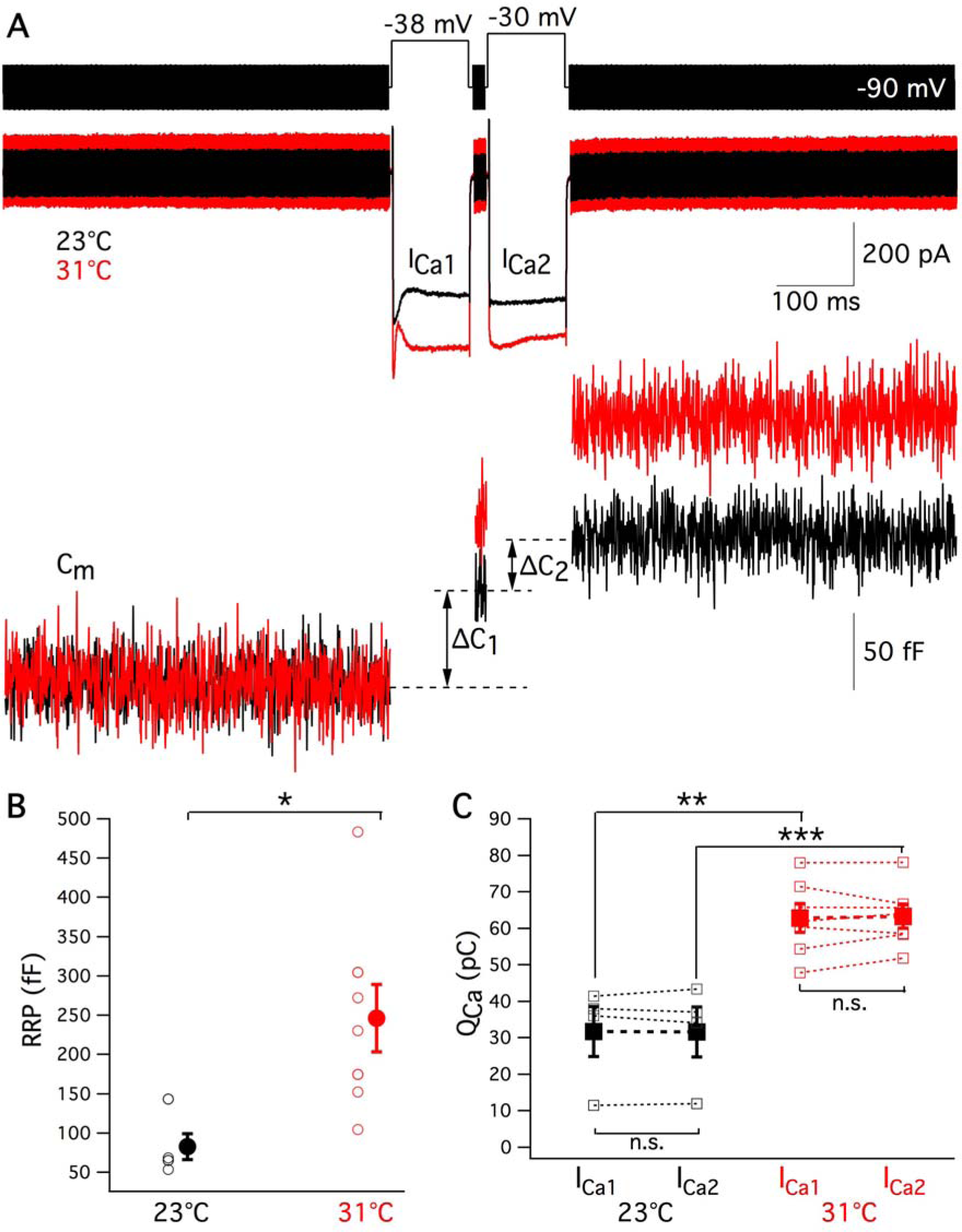
The readily releasable pool of vesicles is larger at high temperature. (A) The paired pulse protocol used to calculate the maximum size of the RRP was composed of two 100 ms pulses with a 25 ms interval. The second pulse was depolarized to −30 mV. The first pulse was adjusted to give the same total amount of Ca^2+^ influx (Q_Ca_) as the second pulse. It was different from cell to cell. In this example, the first depolarizing potential was −38 mV. The intervals between pulse pairs was 60-90 seconds to allow for full recovery. Ca^2+^ current recorded at 23°C and 31°C are shown in black and red, respectively. ΔC_1_ and ΔC_2_ were capacitance jumps triggered by the first and second depolarization, respectively. (B) The maximum size of the RRP was calculated according to 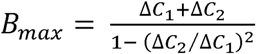. The value of B_max_ increased significantly from about 83 fF (n = 4) at 23°C to 246 fF at 31°C (n = 7, *p* = 0.0351, unpaired *t* test). (C) The Ca^2+^ influx during the first and second depolarizing pulses were 31.8 ± 6.8 pC and 31.7 ± 6.8 pC at room temperature (n = 4, *p* = 0.93), whereas the Ca^2+^ influx during the first and second pulses were 62.8 ± 3.8 pC and 63.3 ± 3.1 pC at high temperature (n = 7, *p* = 0.69). Both the first (*p* = 0.0005, unpaired *t* test) and the second (*p* = 0.0002, unpaired *t* test) pulses thus triggered more Ca^2+^ influx at high temperature.

This equation from Gillis et al. (1996) is derived based on the assumption that the same fraction of the pool is released with each pulse. If there is a larger fraction of release during the second pulse (eg, due to residual Ca^2+^ from the first pulse) or significant numbers of vesicles have been refilled during the second pulse, then B_max_ will overestimate the actual initial pool size. Therefore, the actual pool size lies between ΔC_1_ and B_max_. In the example of Fig. 10A, the B_max_ increased from 143 pF at 23°C to 175 pF at 31°C. On average the B_max_ increased from 83 ± 20 fF (n = 4) at room temperature to 246 ± 47 fF at high temperature (n = 7, *p* = 0.0351, unpaired *t* test; Fig. 10B). These results showing an increase in RRP size with increasing temperature are similar to those reported for chromaffin cell exocytosis (Dinkelacker et al., 2000). In addition, the paired pulse ratio ΔC_2_/ΔC_1_ did not change significantly upon temperature shift: 0.46 ± 0.19 at room temperature (n = 4) vs. 0.69 ± 0.05 at high temperature (n = 7, p = 0.18; unpaired *t* test). This suggests that release probability does not change with temperature shifts.

### Ultrafast Ca^2+^ currents and short synaptic delays at high temperature

*In vivo* recordings of the auditory nerve fibers of two different species of frogs, *Hyla regilla* and *Eleutherodactylus coqui*, showed that the latencies of the first spike after the onset of a sound stimulus decreased when increasing temperature from 12°C to 25°C (Stiebler and Narins, 1990). To determine the temperature dependence of the synaptic delay at the hair cell afferent synapse we performed paired recordings. We depolarized hair cells with two 20-ms depolarizing steps from −90 mV to −30 mV with 50-ms intervals (Fig. 11A) and recorded evoked EPSCs mediated by activation of AMPA receptors in the afferent fiber. After depolarization, Ca^2+^ ions entered into hair cells and triggered glutamate release (Fig. 11A). Due to EPSC rundown in some paired recordings, we only included the first trial from each pair in our analysis. Therefore, the data obtained at room and high temperatures were from two different groups of pairs. The average of EPSCs recorded at 22°C (n = 3) and 31°C (n = 6) are shown in black and red, respectively (Fig. 11A). Synaptic delay was measured from the onset of the pulse until the EPSC reached 10% of its maximum (Fig. 11C). The synaptic delay of the 1^st^ EPSC (Fig. 11C) decreased from 2.03 ± 0.16 ms at room temperature (n = 3) to 1.28 ± 0.15 ms at high temperature (n = 6, *p* = 0.0183, unpaired *t* test; Fig. 11E). The synaptic delay of the 2^nd^ EPSC (Fig. 11C) shortened from 1.26 ± 0.10 ms at room temperature (n = 3) to 0.72 ± 0.05 ms at high temperature (n = 6, *p* = 0.001, unpaired *t* test; Fig. 11E). The 1^st^ synaptic delay was longer than the 2^nd^ one under both room (n = 3, *p* = 0.008) and high (n = 6, *p* = 0.0206) temperature (Fig. 11 D and E). The synaptic delay we measured from adult frog hair cell synapses is much shorter than that of the immature rat inner hair cell synapse (Goutman and Glowatzki, 2011).

**Fig. 11.**
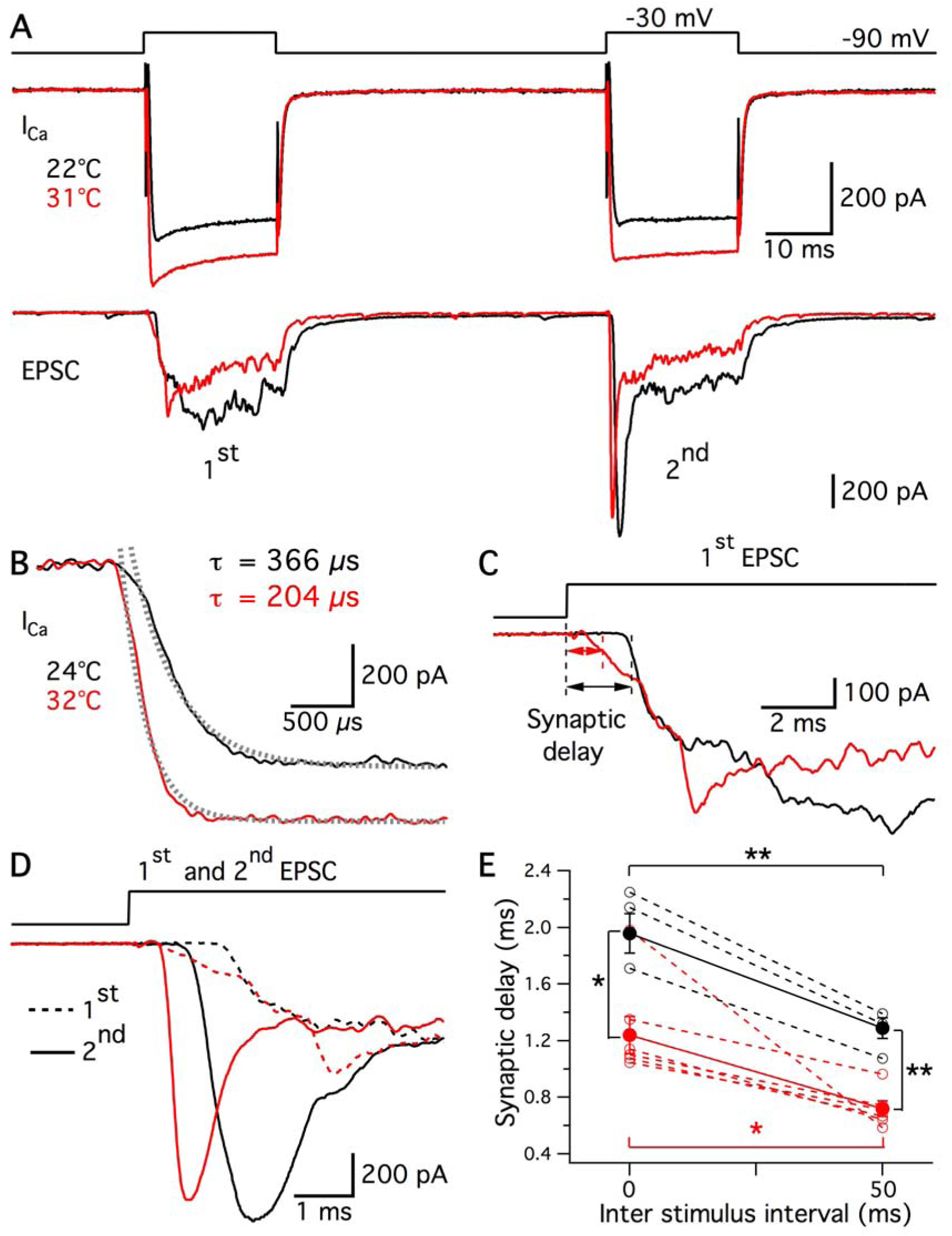
Shorter synaptic delays and faster EPSCs at high temperature. (A) Two 25-ms depolarizing steps from −90 mV to −30 mV with 50 ms intervals were applied to a hair cell. Simultaneously, the post-synaptic afferent fiber was voltage clamped at −90 mV. Average of calcium influx (I_Ca_) elicited by the hair cell stimulus at 22°C (n = 3) and 31°C (n = 6) are shown in black and red, respectively. Synaptic vesicle glutamate release from the hair cell triggered a multiquantal EPSC in the afferent fiber. Black and red traces represent the averaged EPSCs at 22°C (n = 3) and 31°C (n = 6), respectively. Due to significant rundown, we only analyzed the first trial of each pair. Therefore, I_Ca_ and EPSC responses at room and high temperature were obtained from two different groups of pairs. (B) Representative Ca^2+^ currents from another hair cell were triggered by a 20-ms depolarization from −90 mV to −30 mV at 24°C (black) and 32°C (red) using a P/4 protocol for leak subtraction. The activation time constant was derived from the single exponential fit after the capacitive transient (dash lines). In this example, time constants of activation were 366 μs at 24°C (black) and 204 μs at 32°C (red). (C) The rising phase of the first EPSC is shown on an expanded time scale. Synaptic delay was defined as the time between the start of stimulation and to when the EPSC reached 10% of the maximum. (D) Overlay of the first (dashed lines) and the second (solid lines) EPSCs. Note the large synaptic facilitation of the second EPSCs. (E) The synaptic delay of both the 1^st^ (*p* = 0.0183, unpaired *t* test) and 2^nd^ EPSCs (*p* = 0.001, unpaired *t* test) decreased at 31°C. The 1^st^ synaptic delay was longer than the 2^nd^ one under both 22°C (n = 3, *p* = 0.008) and 31°C (n = 6, *p* = 0.0206).

Shorter synaptic delay is partially due to faster activation of Ca^2+^ current at high temperature. In order to determine the effect of temperature on the activation of I_Ca_, we employed P/4 leak subtraction to eliminate the capacitive transient. The Ca^2+^ current was evoked by a 20-ms depolarizing step from −90 mV to −30 mV (Fig. 11B). By fitting the activation phase of the Ca^2+^ current with a single exponential function (Fig. 11B, dashed lines), we calculated the time constant of activation: 377 ± 30 μs at 23.9 ± 0.1°C and 219 ± 19 μs at 31.5 ± 0.3°C (*p* < 0.0001, n = 8). The corresponding Q_10_ was 2.2 ± 0.2 (n = 8), which is similar to the Q_10_ of L-type Ca^2+^ channels activation in chick sensory neurons (Acerbo and Nobile, 1994), but higher than the Q_10_ of gerbil inner hair cells (Johnson and Marcotti, 2008). The results indicate that high temperature speeds up the activation of Ca^2+^ channels and thus contributes to shortening the synaptic delay.

Paired-pulse ratio calculated from the charge transfer of EPSCs (EPSC_2_/EPSC_1_) were 1.00 ± 0.05 at room temperature (n = 3) and 0.92 ± 0.02 at high temperature (n = 6, *p* = 0.25, unpaired *t* test). Together with the paired pulse ratio ΔC_2_/ΔC_1_, this suggests that high temperature does not affect overall release probability. However, high temperature improves temporal precision by shortening the synaptic delay and under both temperatures hair cell synapses exhibit pronounced short-term facilitation (see Cho et al., 2011; Goutman and Glowatzki, 2011). Interestingly, these results differ greatly from those found in a conventional active zone synapse, the calyx of Held synapse, where synaptic delay of the second pulse was longer and EPSCs exhibit pronounced short-term depression (Wu and Borst, 1999).

## Discussion

### Temperature-dependent changes in EPSC, EPSP and spike rates

Our *in vitro* whole-cell recordings of afferent fibers show that the frequencies of spontaneous EPSCs and EPSPs increased at higher temperatures. This should produce an increase in action potential spikes at higher temperatures. Indeed, afferent fiber spike rates mostly increased at higher temperatures (Fig. 1). *In vivo* single auditory nerve fiber recordings also show that spontaneous spike rates in the bullfrog amphibian papilla fibers increase at higher body temperatures (van Dijk et al., 1990). However, about 30% of our recordings of afferent fibers decreased their spike rate at higher temperatures. We do not know the reasons for this heterogeneity. The underlying mechanisms that trigger spikes in the different afferent fibers require additional studies to explain the wide range of membrane input resistance (Fig. 4B). Interestingly, mammalian auditory nerve fibers also exhibit a heterogeneity in spike rates, gene expression and membrane properties (Rutherford et al., 2012; Heil and Peterson, 2017; Shrestha et al., 2018).

What are the mechanisms that produce an increase in EPSC frequency? One obvious possibility we explored was a depolarization of the hair cell V_rest_. Surprisingly, during whole-cell current clamp recordings the V_rest_ of hair cells hyperpolarized significantly (Fig. 5A). V_rest_ also hyperpolarizes in snail neurons at higher temperatures due to increased Na^+^/K^+^-ATPase activity (Gorman and Marmor, 1970). However, perforated patch recordings, which do not disturb the intracellular milieu, showed no significant change. The V_rest_ we observed in perforated patch mode was about −63 mV (Fig. 5A). Previous, sharp conventional electrode recordings of V_rest_ of turtle and frog hair cells reported values around −55 mV (Crawford and Fettiplace, 1980; Pitchford and Ashmore, 1987). At these membrane potentials a small inward Ca^2+^ current is induced (Fig. 5C). This resting Ca^2+^ current increases at higher temperatures, which probably increases release probability. The frequency of spontaneous mEPSCs also increases at the calyx of Held synapse (Kushmerick et al., 2006; Postlethwaite et al., 2007) and hippocampal synapses (Kim and Connors, 2012) at high temperature. However, unlike bullfrog hair cells, V_rest_ of the calyx of Held nerve terminal depolarizes when temperature increases from 24°C to 32°C (Kim and von Gersdorff, 2012). Finally, we note that our amphibian papilla hair cells are bathed in high external Ca^2+^, which blocks transduction channels, and recently, it was found that sudden heat rises can soften gating springs and open mechanotransduction channels on the stereocilia bundles of bullfrog sacculus hair cells (Azimzadeh et al., 2018). Therefore, we may be underestimating the physiological effects of temperature shifts on the hair cell and V_rest_ may be more depolarized under *in vivo* conditions.

We found out that the average amplitude of the spontaneous EPSCs recorded from bullfrog auditory afferent fibers did not change at higher temperature (Fig. 2 and Table 2). This differs from observations made in mammalian synapses where elevating temperature increases spontaneous mEPSCs amplitude (Veruki et al., 2003; Kushmerick et al., 2006; Postlethwaite et al., 2007). Interestingly, we previously observed that a decrease in temperature from room temperature to 15°C resulted in a decrease in EPSC amplitude (Li et al., 2009). Spontaneous EPSC amplitudes may thus be saturated at room temperature. The average EPSP amplitude also did not change with temperature (Table 2). However, in 13 fibers were both EPSCs and EPSPs were recorded, EPSCs remained the same (*p* = 0.45), whereas EPSP amplitude decreased (*p* = 0.049, n = 13) at high temperature. This could result from a decrease in afferent fiber R_in_ at high temperature. Both EPSC and EPSP events increased in frequency at high temperature (Table 2). This can lead to greater summation of EPSPs making them more likely to reach action potential threshold. However, EPSP decay was also significantly faster at higher temperatures (Fig. 2A3 and 2B3), which tends to reduce summation of non-coincident EPSPs. Therefore, it is possible that higher temperatures may reduce spike rates, as shown in Fig. 1B. Presynaptic and postsynaptic mechanisms may thus explain the heterogeneity in spike frequency changes with temperature seen in Figure 1.

Temperature-dependent changes in spike threshold may also explain the heterogeneity shown in Fig. 1. Thermo-positive fibers fire more spikes spontaneously at high temperature with threshold decreasing by about 5 mV, whereas thermo-negative fibers fire less spontaneous spikes at high temperature with threshold increasing by 5 mV (Table 1). Heterogeneity in the types and densities of voltage-gated Na^+^ and K^+^ channels among different fibers may explain these differences (Kim and Rutherford, 2016). Altogether, the temperature-dependent changes of spiking rates is due to presynaptic changes in hair cell exocytosis and postsynaptic changes in afferent fiber excitability.

### Synaptic delay and phase locking at high temperature

Recordings of single frog auditory nerve fibers show that spike onset delay to sound stimulus decreased as temperature increased with Q_10_ values of 1.6-2.5 (Stiebler and Narins, 1990; van Dijk et al., 1990). We performed paired recordings from hair cell to afferent fiber and revealed a shortening of synaptic delay at high temperature (Fig. 11E). This shorter synaptic delay was due in part to a faster activation kinetics of Ca^2+^ current at high temperature (Fig. 11B). In addition, the time constant of lipid membrane fusion is inversely correlated to temperature (Schotten et al., 2015), which may also contribute to the shorter synaptic delay at high temperature. A shorter synaptic delay will improve the spike phase-locking of the fibers because evoked EPSPs and spikes will be more phase-locked to the hair cell Ca^2+^ current activation. This may explain why *in vivo* recordings from single auditory nerve fibers indicate that vector strength increases at higher temperatures (Stiebler and Narins, 1990; van Dijk et al., 1990).

Higher temperatures and synaptic facilitation tend to synchronize depolarization evoked multivesicular release (Fig. 11). Synaptic facilitation revealed by paired-pulse depolarization also reduced synaptic delays in both bullfrog hair cell synapses (Cho et al., 2011) and rat inner hair cell synapses (synaptic delay was 1.5 ms at room temperature; Goutman and Glowatzki, 2011). Likewise, we measured here synaptic delays of 1.3 ms at 23°C and 0.7 ms at 31°C (Fig. 11E).

### Electrical resonance at high temperature

The lowering of R_in_ at high temperature reduces the membrane time constant, allowing hair cells to better follow rapid membrane potential changes (Fig. 6). The intrinsic electrical resonant frequency of hair cells increased with temperature (Fig. 7), as expected from an increase in Ca^2+^ and K^+^ current amplitudes (Fig. 5B). We observed a large change in average resonant frequency from 320 Hz to 530 Hz (Fig. 7D). These changes increase the dynamic range of frequency tuning of amphibian papilla hair cells and their connected afferent fibers. Indeed, *in vivo* recordings of frog auditory nerve fibers show higher characteristic frequencies (tuning curve tip or best frequency) at higher temperature (Stiebler and Narins, 1990).

### Exocytosis efficiency and vesicle pool size at high temperature

Smaller R_in_ will inevitably result in smaller membrane potential changes (V_m_ = R_in_ × I) induced by the opening of mechanosensitive channels in the stereocilia. This predicted loss of sensitivity to sound signals is not observed *in vivo*. In fact, the contrary is observed: the threshold for sound triggered spikes is reduced at high temperature (Stiebler and Narins, 1990). How can we explain this increase in fiber sensitivity?

First, we found that the steepness of the Ca^2+^ current IV curve (Fig. 5C) increased at high temperature making Ca^2+^ influx more sensitive to membrane potential fluctuations around V_rest_. The enlarged Ca^2+^ current could result from an increase in L-type Ca^2+^ channel open probability (P_o_) and/or conductance (Lux and Brown, 1984; Klöckner et al., 1990; Acerbo and Nobile, 1994; Peloquin et al., 2008). The Q_10_ of single L-type channel conductance in smooth muscle cells and retina is 1.6 and 3.0, respectively (Klöckner et al., 1990; Peloquin et al., 2008). Second, at hair cell synapses a cluster of L-type Ca^2+^ channels is located within nanometers from docked vesicles under the synaptic ribbon (Brandt et al., 2005; Nouvian et al., 2006; Graydon et al., 2011; Kim et al., 2013). Fast activation of this cluster of Ca^2+^ channels produces a highly synchronous and fast multiquantal EPSC (Fig. 11D; Grabner et al., 2016), which may help to explain the short latency and low jitter of the sound evoked first-spike in the auditory nerve (Wittig and Parsons, 2008).

We also observed an increase in exocytosis efficiency, which indicates the exocytosis increase is not proportional to the increase of Ca^2+^ current. A nonlinear power law dependence of exocytosis on free Ca^2+^ ion concentration could potentially explain our results of a high Q_10_ = 2.4 for exocytosis elicited by 20-ms depolarizing pulses (Cho and von Gersdorff, 2012; Quiñones et al., 2012). However, a linear dependence of exocytosis on Ca^2+^ current charge has been found for mid-frequency tuned bullfrog hair cells (Keen and Hudspeth, 2006; Cho et al., 2011). Thus, vesicle fusion efficiency must be intrinsically higher at high temperature, as shown in Fig. 9C. Besides Ca^2+^ influx through L-type Ca^2+^ channels in the plasma membrane, Ca^2+^-induced Ca^2+^ release from intracellular stores can also be involved in nonlinear vesicle recruitment at auditory hair cells (Castellano-Muñoz et al., 2016).

Release probability increases at high temperature for some central synapses (Hardingham and Larkman, 1998; Volgushev et al., 2004), although we did not observe changes in release probability using a paired-pulse stimulation protocol (Fig. 10 and 11). However, we did observe a significant increase in the readily releasable pool (RRP) size at high temperature (Fig. 10). In addition to facilitating spontaneous vesicle fusion, temperature also facilitates sustained exocytosis by accelerating vesicle replenishment of the RRP (Pyott and Rosenmund, 2002; Kushmerick et al., 2006). Moreover, synaptic vesicle endocytosis is also highly temperature sensitive (Fernández-Alfonso and Ryan, 2004; Micheva and Smith, 2005; Renden and von Gersdorff, 2007; Delvendahl et al., 2016). The kinetics of endocytosis may thus be rate limiting for prolonged exocytosis and requires further studies at hair cell synapses.

## Acknowledgements

This research was supported by the National Institutes of Health grant to HvG (NIDCD DC004274). We thank Jutta Engel for conversations that inspired this study, Karina Leal, Geng-Lin Li, and Owen Gross for data analysis assistance, and Chad Grabner for very insightful discussions.

